# BRIDGE: inter-tissue imputation with contrastive learning improves recovery of donor-specific gene expression

**DOI:** 10.64898/2026.11.24.768988

**Authors:** Jingbo Yang, Ronnie Y. Li, Zhaohui S. Qin

## Abstract

Multi-tissue transcriptome resources are central to understanding tissue-specific regulation and inter-individual variation. Realistically, most tissues are measured in only a subset of donors, especially for less accessible tissues such as brain and heart. Accurately imputing gene expression values for such tissues, while accounting for individual heterogeneity, remains a daunting challenge. Here, we present BRIDGE (Body-wide Representation via Inter-tissue Decoding of Gene Expression), a lightweight framework that imputes donor-specific and tissue-specific expression measures using contrastive learning. When tested on real-world expression data from the Genotype-Tissue Expression (GTEx) consortium, BRIDGE recovered donor-specific variation more accurately than existing state-of-the-art methods while improving overall accuracy, with the largest advantage found in the most inaccessible tissues. We provide the BRIDGE-imputed completed GTEx v11 donor × tissue × gene expression tensor as a resource to the broader research community.

**Key points:**

1. Existing tissue-expression imputation methods are judged on whole-profile accuracy, which the tissue mean alone nearly saturates, so we evaluate the recovery of donor-specific deviations instead.
2. We present BRIDGE, which combines a donor-level average expert and a contrastively trained sample matching expert through an adaptive mixer, using only tissue identity and gene expression.
3. In GTEx hide-one imputation, BRIDGE more than doubled the residual Pearson correlation of HYFA and retained its advantage when only four accessible tissues were observed.
4. BRIDGE-imputed data better preserved gene-age associations and Hallmark pathway signals than HYFA, with the largest gains at high missingness, although imputation still shrinks effect sizes.
5. BRIDGE trains quickly, improves with more donors, and is released as a measured-plus-imputed GTEx v11 resource covering 946 donors.

## Introduction

Gene expression varies across tissues and individuals, reflecting the complex interplay between tissue-specific regulatory programs, genetic variation, and environmental influences. Characterizing these patterns across multiple tissues is essential for understanding shared and tissue-specific gene regulation, inter-individual variation, and the molecular mechanisms connecting regulatory variation to complex traits and diseases^1-4^. Large-scale resources such as the Genotype-Tissue Expression (GTEx) consortium have enabled these investigations by providing matched transcriptomic profiles across diverse tissues collected from the same donors^1,2^. However, broad tissue coverage at the cohort level does not translate into complete multi-tissue profiles for individual donors, as most tissues are collected from only a subset of individuals^5^. This limitation is particularly pronounced for difficult-to-access tissues, such as those from the brain and heart. Accurately imputing expression in unobserved tissues is therefore essential for maximizing the utility of multi-tissue transcriptomic resources and enabling more comprehensive characterization of individual-level gene expression across tissues.

A central challenge in multi-tissue expression imputation is recovering donor-specific variation rather than simply reproducing average tissue expression patterns. This distinction is important because inter-individual differences in gene expression may reflect biologically meaningful regulatory variation that is obscured when predictions are dominated by tissue-level expression patterns. Conventional evaluation metrics provide limited evidence of a method’s ability to recover this variation. For example, assigning each missing sample the mean expression profile of its target tissue yields an almost perfect Pearson correlation, largely because broad differences in mean expression across genes dominate this metric. Thus, high sample-wise accuracy can conceal a failure to recover differences between individuals. To address this limitation, we focus on residual expression, defined as the deviation of a donor’s measured expression profile from the corresponding tissue mean. By evaluating deviations from average tissue expression, residual-based metrics provide a more direct assessment of whether an imputation method preserves donor-specific expression variation. Accordingly, accurate recovery of residual expression is a central objective of our approach.

Previous approaches to tissue-expression imputation have leveraged statistical matching, surrogate-tissue measurements, genetic predictors, and shared regulatory information across tissues. Predictive mean matching (PMM) imputes missing values using observed cases with similar model-based predictions^6,7^. Surrogate-tissue approaches use expression measured in accessible tissues, most commonly whole blood, to infer expression in less accessible tissues^8,9^. However, their utility is constrained by tissue- and cell-type-specific regulation, which can limit the transferability of expression patterns across tissues^2^. Genotype-based models predict tissue-specific expression from regulatory variants^10–12^, providing a complementary strategy that leverages genetic information but requires individual-level genotypes and primarily captures genetically regulated components of expression. Other approaches, including mixed-effects and multi-tissue methods such as Meta-Tissue, leverage shared genetic effects across tissues to improve the identification of tissue-specific regulatory associations^13,14^. TEEBoT predicts target-tissue expression by projecting whole-blood expression and splicing profiles onto low-dimensional principal components and applying tissue-specific linear regression^15^. Related approaches have extended surrogate-based prediction using Bayesian or multi-modal regression^16-18^, while multi-task and deep learning models have sought to capture more complex relationships across tissues and predict tissue-specific expression profiles^19,20^. Collectively, these studies demonstrate the feasibility of tissue-expression imputation and the value of leveraging information across tissues and individuals. Nevertheless, accurately recovering donor-specific expression variation from incomplete multi-tissue profiles remains challenging. Approaches that rely on a limited number of surrogate tissues may not fully exploit information distributed across multiple observed tissues, while methods based on statistical matching or genetic prediction may not directly leverage similarities between donors to recover individual-specific expression patterns. These limitations motivate methods that integrate within-donor information across tissues with between-donor similarities to improve the recovery of expression in unobserved tissues.

More recently, Hypergraph Factorization (HYFA) introduced a framework for multi-tissue expression imputation that represents the observed tissue-expression profiles of each donor, together with age and sex, in a hypergraph^5^. Through an encoder, message-passing network, and decoder, HYFA learns factorized donor, tissue, and metagene representations for target-tissue prediction. This formulation accommodates variable sets of observed tissues, captures nonlinear cross-tissue dependencies, and does not require individual-level genotype data. Although HYFA represents an important advance in genotype-independent multi-tissue imputation, its learned representations do not explicitly identify similar donors in the reference population whose expression profiles may provide informative analogies for individual predictions. Similarly, its tissue embeddings and metagene enrichments provide global summaries of biological relationships but do not directly reveal which observed tissues or reference donors contribute to a particular imputation. Explicitly leveraging donor-level similarities could therefore provide a complementary means of recovering individual-specific expression variation while offering greater transparency into the information underlying each prediction.

Here, we introduce BRIDGE (Body-wide Representation via Inter-tissue Decoding of Gene Expression), a lightweight framework that comprises two complementary prediction experts: a donor-level average expert and a sample matching expert. Each expert produces a complete gene-expression estimate, and a sample-adaptive mixer combines their predictions using learned weights. First, a donor-level average expert leverages expression relationships across tissues within the same individual to inform predictions. Second, a contrastively trained sample-matching expert identifies similar donors in the reference population and uses their expression profiles to recover missing values through between-donor analogy. These complementary experts capture within-donor and between-donor information, respectively, allowing BRIDGE to exploit both the observed expression profile of an individual and informative similarities across the reference population. An adaptive mixer integrates their predictions according to the target tissue and the number of observed tissues available for each donor, allowing the model to adjust the relative contribution of each information source under varying degrees of tissue availability. The retrieved neighbors and associated expert weights are directly inspectable, providing transparency into the reference information used for individual predictions.

Using GTEx v11, we demonstrate that BRIDGE improves the recovery of donor-specific expression variation compared with HYFA while maintaining or improving overall expression prediction accuracy. Its advantages are particularly pronounced when only a limited number of accessible tissues are available, addressing an important challenge in the application of multi-tissue imputation to incompletely sampled donors. BRIDGE also achieves matched validation error in fewer training epochs and generally provides more accurate recovery of age-associated expression effects. Finally, we use BRIDGE to generate a completed GTEx v11 expression resource containing both measured and imputed donor-tissue-gene profiles, with the two clearly distinguished to facilitate downstream use. We further demonstrate that the donor-retrieval component continues to improve as additional training donors are incorporated, highlighting the potential value of expanding reference populations for donor-aligned expression imputation.

## Methods

### GTEx data

The Genotype-Tissue Expression (GTEx) consortium has exerted tremendous effort in conducting bulk RNA-sequencing profiling across a broad range of non-diseased human tissues, with the goal of elucidating tissue-specific gene expression and regulation^1,2^. Here, we used processed gene expression data in transcripts per million (TPM) from GTEx version 11, which comprised 19,788 tissue profiles collected from 946 donors and 68 different tissues. Each donor contributed 20.9 tissues on average.

### Study design

All 68 tissues appearing in the GTEx v11 dataset were retained in the present study, and 945 donors with at least two available tissues were included. To evaluate performance, we divided 945 donors into three subsets in a 7:1:2 ratio: 662 in training, 95 in validation, and 188 in testing. Three tissues (2–8 donors each) were present only in the training set, so performance on them could not be appropriately evaluated; thus, missing rates are reported for the remaining 65 tissues, averaging 67.8% (range 12.8%–99.2%; **Supplementary Figure 1, Supplementary Figure 2**).

### Data processing

Expression values were represented as ln(1 + TPM). To remove non-informative genes, we ranked genes by their variance across all tissue samples and retained the top 20,000 genes with the highest variance for all downstream analyses. For tissue *t*, we first calculated its average expression profile ***µ***_*t*_ = (*µ*_*t*,1_, *µ*_*t*,2_, …, *µ*_*t*,20000_) across all training donors with this tissue type. For the tissue sample from donor *i* and tissue *t*, we used profile ***x***_*i,t*_ = (*x*_*i,t*,1_, *x*_*i,t*,2_, …, *x*_*i,t*,20000_), to represent the observed expression profile. We defined residual expression profile as

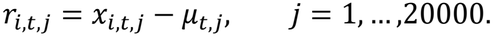

### BRIDGE model

BRIDGE imputes expression in an unobserved tissue by adaptively combining predictions from two complementary experts: (1) the donor-level average expert, which captures donor-specific signals from the donor’s observed tissues, and (2) the sample-matching expert, which retrieves similar tissue-expression profiles from the training cohort. A schematic of the model architecture is provided in **Figure 1**.

**Figure 1.**
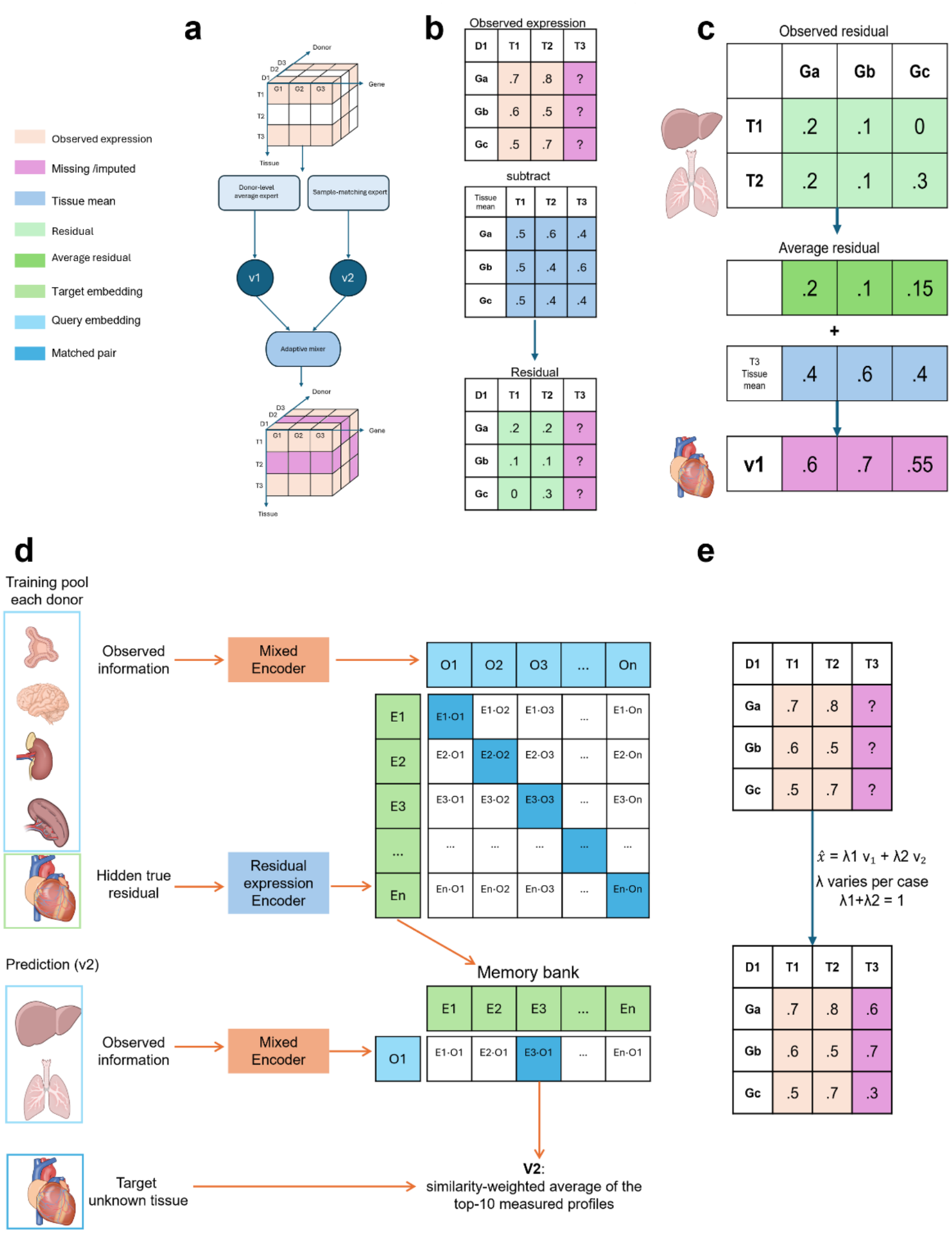
Schematic overview of BRIDGE. (a) The overall BRIDGE workflow. (b) Residual expression is obtained by subtracting the training-derived tissue mean from the observed expression; question marks indicate an unobserved tissue in a donor. (c) The donor-level average expert adds the donor’s average residual across observed tissues to the mean profile of the missing tissue, giving prediction *v*_1_. (d) The sample matching expert retrieves similar measured profiles from training donors and averages the ten closest matches to give prediction *v*_2_. (e) An adaptive mixer combines *v*_1_ and *v*_2_ to fill in the missing tissue. Values are illustrative.

#### Donor-level average expert

The donor-level average expert predicts the target-tissue profile from the donor’s other measured tissues alone. Let *K*_*i,t*_ denote the set of tissues measured for donor *i*, excluding the target tissue *t*. The prediction for gene *j* is the target-tissue mean plus the donor’s average residual across *K*_*i,t*_:

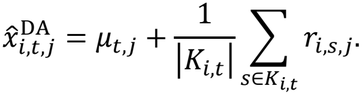

Here *µ*_*t,j*_ is the training-cohort mean of gene *j* in tissue *t*, and *r*_*i,s,j*_ is the residual of donor *i* in source tissue *s* (see **Data processing**). This is analogous to estimating a donor-specific random intercept shared across tissues^13^, and it assumes that donor-specific deviations in expression are partially shared across tissues.

#### Sample matching expert

The sample matching expert retrieves measured training profiles that resemble the evidence available for a donor and uses them to predict the target tissue. It has three parts: an encoder that maps donor evidence and target profiles into a shared space, a contrastive objective that trains that space, and a memory bank from which neighbors are retrieved at prediction time.

#### Query and target representations

Each training record (*i, t*) pairs a donor *i* with a held-out target tissue *t* and uses the donor’s remaining measured tissues, *K*_*i,t*_, as evidence. Records are built by holding out each measured tissue in turn (see Hide-one-tissue imputation).

First, each observed residual r_*i,s*_, *s* ∈ *K*_*i,t*_, is passed through the shared encoder *E* and *L*_2_-normalized:

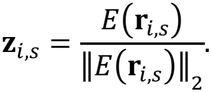

Second, each **z**_*i,s*_ is concatenated (⊕) with the embedding **e**_*s*_ of its source tissue, transformed by *C*, and averaged over the observed tissues to give the donor evidence state:

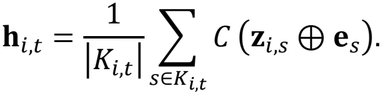

Because the average is taken over a set, **h**_*i,t*_ does not depend on tissue order and accommodates any number of observed tissues^22^.

Third, the evidence state is concatenated with the target-tissue embedding **e**_*t*_, transformed by *Q*, and *L*_2_-normalized to form the query:

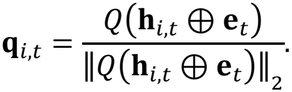

Finally, the held-out target residual r_*i,t*_ is passed through the same encoder *E* to give the target representation:

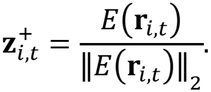

The query summarizes what the donor’s other tissues imply about tissue *t*, whereas 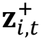 describes what that tissue actually looks like. Because the same encoder is applied to source and target residuals, queries and target representations lie in one space.

The encoder *E* is a multilayer perceptron with one hidden layer of 512 units and a Gaussian Error Linear Unit (GELU) activation that maps the 20,000-gene residual to a 128-dimensional representation. The source-and target-tissue embeddings **e**_*s*_ and **e**_*t*_ are 32-dimensional vectors that are learned, being updated at each training step. The inputs to *C* and *Q* are therefore 160-dimensional (128 + 32), and each returns a 128-dimensional vector, so queries and target representations are both 128-dimensional.

#### Contrastive training

We trained *E, C* and *Q* with a symmetric InfoNCE objective^23,24^, which teaches the model to pair each query with the target representation of its own record. For each training record, the query **q**_*i,t*_ served as the anchor, and the target representation 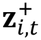 from the same donor and held-out tissue served as the positive. Target representations from all other records in the same minibatch served as in-batch negatives^25^. A minibatch can contain several records from the same donor, and these act as negatives for one another.

For a minibatch of *B* records indexed by *a* and *b*, let record *a* have donor *i*_*a*_ and target tissue *t*_*a*_, and write 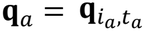 and 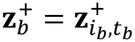. The temperature-scaled similarity was

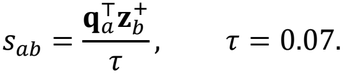

Because both vectors are *L*_2_-normalized, 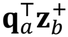 is their cosine similarity. The symmetric InfoNCE loss was

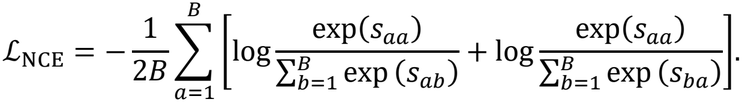

The first term optimized query-to-target matching: each query must identify its own target among the *B* targets in the batch. The second optimized target-to-query matching^26^: each target must identify its own query.

#### Memory bank and retrieval

After contrastive training, each training record *n* was stored as a key–value pair^26,27^. The key **z**_*n*_ is the record’s normalized target-residual representation (its **z**^+^), and the value **x**_*n*_ is its measured target-tissue expression profile in ln(1 + TPM) space, not a residual. Writing *t*_*n*_ for the target tissue of record *n*, the memory bank and its target-tissue subsets are

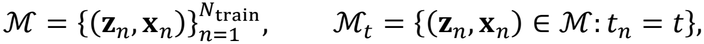

with *N*_train_ = 13,879 in the hide-one benchmark.

Retrieval was not limited to the target tissue and could return other tissues. At prediction time, the query **q**_*i,t*_ was compared by cosine similarity with the keys in ℳ _*t*_ only, and *N*_*k*_(**q**_*i,t*_) denotes the *k* = 10 entries of ℳ _*t*_ with the highest cosine similarity. Their measured expression profiles were averaged using softmax-normalized similarity weights^27,28^, with no temperature applied:

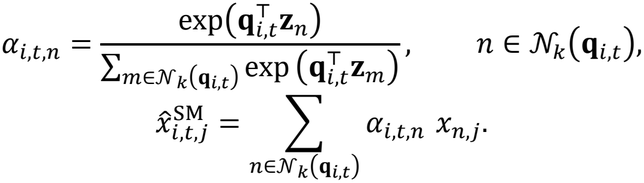

Because every retrieved value is a profile of the target tissue, and the values are expression profiles rather than residuals, the prediction is on the same scale as the donor-level average prediction.

#### Adaptive prediction mixer

An adaptive mixer^21^ determined how much each expert contributed to each prediction. It took the target-tissue identity and the number of tissues observed for the donor, scaled as

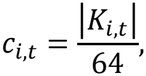

Here 64 is the largest possible number of observed source tissues because one tissue is hidden as the target in each iteration (65 − 1 = 64). The mixer produced two softmax-normalized weights:

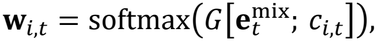

where *G* is the mixer network and 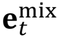 is the mixer’s target-tissue embedding. The weights combine the donor-level average and sample matching predictions:

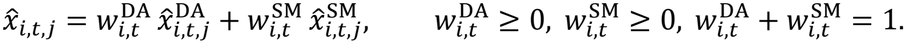

The two experts were held fixed while the mixer was trained against the measured target-expression profiles using Smooth L1 loss^29^. During mixer training, the target record being predicted was excluded from its own sample matching candidates.

### Hide-one-tissue imputation

Primary performance was assessed in a hide-one setting, which reconstructs one measured tissue from the same donor’s remaining tissues^5^. Each measured tissue was held out in turn as the target, giving 13,879 training, 1,986 validation and 3,922 test records within the respective donor-level partitions (662, 95 and 188 donors; see Study design), all limited to the 65 tissues present in the training set. All methods and ablations were evaluated on the same 3,922 test records across 20,000 genes.

### Accessible tissue imputation

We also considered a practical setting in which donor evidence was limited to four comparatively accessible tissues: Whole Blood, Skin – Sun Exposed (Lower leg), Skin – Not Sun Exposed (Suprapubic) and Adipose – Subcutaneous^5^. Test donors with all four source tissues measured were retained, and each of their remaining measured tissues was held out in turn as the target. Target tissues represented by fewer than 25 test records were excluded^5^, leaving 1,270 records across 28 target tissues. The GTEx-trained BRIDGE and HYFA models were applied directly, without retraining or fine-tuning. We additionally evaluated the age- and sex-adjusted TEEBoT implementation^15^ used in the HYFA notebook^5^, which applies a target-specific, centered 30-component singular-value decomposition followed by multi-output ordinary least-squares regression fitted on training donors only.

### Competing methods and ablations

BRIDGE was benchmarked against HYFA and predictive mean matching (PMM)^6,7^ using identical target records and observed inputs. HYFA was retrained on the same 20,000-gene expression matrix and donor partition, using the architecture and training objective from its official implementation. The donor-level average and sample matching experts were also evaluated separately. A demographically similar control replaced learned retrieval with the ten nearest training donors selected by age and sex, and it was tested both alone and in combination with the donor-level average expert.

Two ablations tested the role of tissue identity. The no-target-identity variant retained source-tissue identities but removed target-tissue identity from the retrieval query and the mixer. The no-any-identity variant additionally removed source-tissue identities from the retrieval query. Both retained the training-derived mean profile of the target tissue used by the donor-level average expert. All comparators and ablations were evaluated on the same 3,922 hide-one test records and 20,000 genes.

### Performance metrics

Residual Pearson correlation was the primary measure of donor-specific expression recovery. For each test record (*i, t*), the training-derived target-tissue mean was subtracted from both the measured and predicted profiles, and the Pearson correlation was calculated across the 20,000 genes:

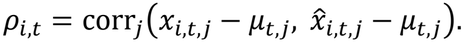

The record-level correlations *ρ*_*i,t*_ were then averaged across the 3,922 test records. Absolute prediction error was measured by mean absolute error (MAE) in ln(1 + TPM) space. Residual gene Pearson, RMSE, sample-wise Pearson correlation and gene-wise Pearson correlation were also reported^5^; full definitions are given in the **Supplementary Materials**.

### Recovery of gene-wise age associations

We assessed whether BRIDGE-reconstructed expression preserved gene-wise associations with donor age^13,30,31^. The analysis included 3,862 hide-one records from 49 test-set tissues that had complete public age and sex information and at least 25 records per tissue. Within each tissue, 20%, 40%, 60% or 80% of the measured profiles were masked, using five fixed age- and sex-stratified masks at each missingness level. The same masks were applied across all analysis arms.

The Oracle arm used all measured profiles, whereas the observed-only arm excluded masked donors. In the mean-fill arm, each masked profile was replaced with gene-wise means estimated from the unmasked donors^32^. In the three imputation arms, masked profiles were replaced with predictions from BRIDGE, HYFA or PMM. Within each tissue and analysis arm, the following linear model was fitted separately for each gene *g*:

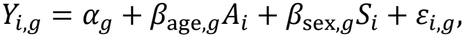

where *Y*_*i,g*_ is the measured or reconstructed expression of gene *g* for donor *i, α*_*g*_ is the gene-specific intercept, *β*_age,*g*_ and *β*_sex,*g*_ are the age and sex coefficients, *A*_*i*_ is the standardized midpoint of the donor’s public age interval, *S*_*i*_ is donor sex, and *ε*_*i,g*_ is the error term.

Recovery was evaluated primarily by the Pearson correlation, across the 20,000 genes, between the estimated age coefficients from each analysis arm and the corresponding Oracle coefficients. Secondary measures were age-coefficient MAE, regression slope, sign agreement, and recovery of the genes with the largest absolute Oracle age coefficients.

### Hallmark gene-set analysis

Residual recovery was first calculated for each of the 20,000 evaluated genes across the hide-one test records. The evaluated genes were then intersected with the 50 human MSigDB Hallmark gene sets^33,34^, and recovery for each Hallmark set was summarized as the mean residual gene Pearson correlation across the genes present in both the set and the evaluated gene panel.

To compare BRIDGE with HYFA, gene-level improvement was calculated across the same hide-one test records as the HYFA MAE minus the BRIDGE MAE. Genes were ranked across the complete evaluable gene panel, and the top 10% most improved genes were defined globally. Enrichment of these genes within each Hallmark set was assessed with a one-sided Fisher’s exact test^35^, followed by Benjamini–Hochberg correction^36^ across the 50 Hallmark sets.

### Training convergence and data scaling

BRIDGE and HYFA were trained for up to 200 epochs on the same donor partition, with validation MAE evaluated after each epoch. Convergence was assessed using the best validation epoch and the first epoch reaching each predefined MAE threshold. For the data-scaling analysis, BRIDGE was independently retrained from seed 42 using 10%, 25%, 50%, 75% or 100% of the training donors, with the validation partition and the 3,922 hide-one test records held fixed.

### Production of the BRIDGE-completed GTEx resource

BRIDGE was retrained on all eligible measured GTEx v11 profiles, using a fixed 200-epoch memory encoder (the contrastively trained *E, C* and *Q* of the sample matching expert) and a 50-epoch adaptive mixer. Tissue eligibility was determined using prespecified minimum-support criteria. The retrieval memory contained only originally measured profiles, and all records from the query donor were excluded during retrieval. Each missing profile was predicted exclusively from that donor’s originally measured tissues; imputed profiles were never reused as model inputs. Quality-control checks tested for duplicate predictions, measured–imputed overlap, query-donor retrieval leakage and non-finite output values.

## Results

### BRIDGE improves donor-specific residual recovery in GTEx hide-one imputation

We first evaluated BRIDGE in the GTEx hide-one setting, in which a measured tissue was withheld from a test donor and predicted from that donor’s remaining tissues, with each measured tissue held out in turn. Across 3,922 held-out records and 20,000 genes, BRIDGE achieved a mean residual Pearson correlation of 0.434, compared with 0.237 for HYFA and −0.012 for PMM, and it had the lowest MAE (0.246, versus 0.259 and 0.797; **Table 1**). BRIDGE also had the highest sample-wise Pearson correlation (0.975), but the target-tissue mean alone reached 0.969, so this metric separates the methods far less than residual correlation does. In the 13 brain tissues, BRIDGE had the highest residual Pearson correlation in every tissue, with pooled means of 0.419, 0.231 and −0.030 for BRIDGE, HYFA and PMM, respectively (**Figure 2a**). Results for all 65 tissues can be found in **Supplementary Data** and **Supplementary Figure 3**, and results for the brain tissues is in **Supplementary Figure 4**.

**Table 1.** Comparison of BRIDGE with HYFA, PMM, TEEBoT and internal model ablations in the GTEx hide-one and accessible-tissue tasks.

| **Task** | **Group** | **Method** | **Residual Pearson** | **Gene residual Pearson** | **MAE** | **RMSE** | **Sample Pearson** | **Gene Pearson** |
|---|---|---|---|---|---|---|---|---|
| Hide-one | Full model | BRIDGE | 0.434 | 0.334 | 0.246 | 0.370 | 0.975 | 0.876 |
|  | Tissue-identity ablation | BRIDGE without target-tissue identity | 0.272 | 0.298 | 0.282 | 0.407 | 0.970 | 0.860 |
|  | Tissue-identity ablation | BRIDGE without any tissue identity | 0.271 | 0.294 | 0.282 | 0.407 | 0.970 | 0.860 |
|  | BRIDGE component | Donor –level average expert | 0.361 | 0.309 | 0.262 | 0.387 | 0.973 | 0.863 |
|  | BRIDGE component | Sample matching expert | 0.318 | 0.202 | 0.263 | 0.400 | 0.971 | 0.853 |
|  | Demographic control | Age/sex control mixer | 0.367 | 0.306 | 0.258 | 0.383 | 0.973 | 0.866 |
|  | Demographic control | Age/sex top-10 only | 0.009 | 0.041 | 0.280 | 0.421 | 0.968 | 0.834 |
|  | Baseline | HYFA | 0.237 | 0.278 | 0.259 | 0.418 | 0.967 | 0.849 |
|  | Baseline | PMM | -0.012 | -0.003 | 0.797 | 1.176 | 0.735 | 0.008 |
|  | Baseline | Target-tissue mean | NA | NA | 0.272 | 0.408 | 0.969 | 0.846 |
| Accessible tissues | Full model | BRIDGE | 0.363 | 0.283 | 0.250 | 0.378 | 0.973 | 0.857 |
|  | Baseline | TEEBoT | 0.349 | 0.255 | 0.250 | 0.383 | 0.972 | 0.853 |
|  | Baseline | HYFA | -0.010 | 0.027 | 0.301 | 0.488 | 0.952 | 0.782 |
|  | Baseline | PMM | -0.018 | -0.016 | 0.755 | 1.124 | 0.767 | -0.009 |

### BRIDGE retains imputation accuracy using accessible source tissues

We next evaluated BRIDGE with the four accessible tissues as the only observed inputs. BRIDGE achieved the highest residual Pearson correlation (0.363), compared with 0.349 for TEEBoT, −0.010 for HYFA and −0.018 for PMM. BRIDGE and TEEBoT had comparable MAE (both 0.250 after rounding), whereas HYFA and PMM had higher errors (0.301 and 0.755, respectively; **Table 1**). Across the 28 target tissues, BRIDGE consistently outperformed HYFA and PMM, and it retained stronger donor-specific residual recovery than TEEBoT despite their similar absolute errors (**Figure 2b, Supplementary Figure 4**, and **Supplementary Data**).

**Figure 2.**
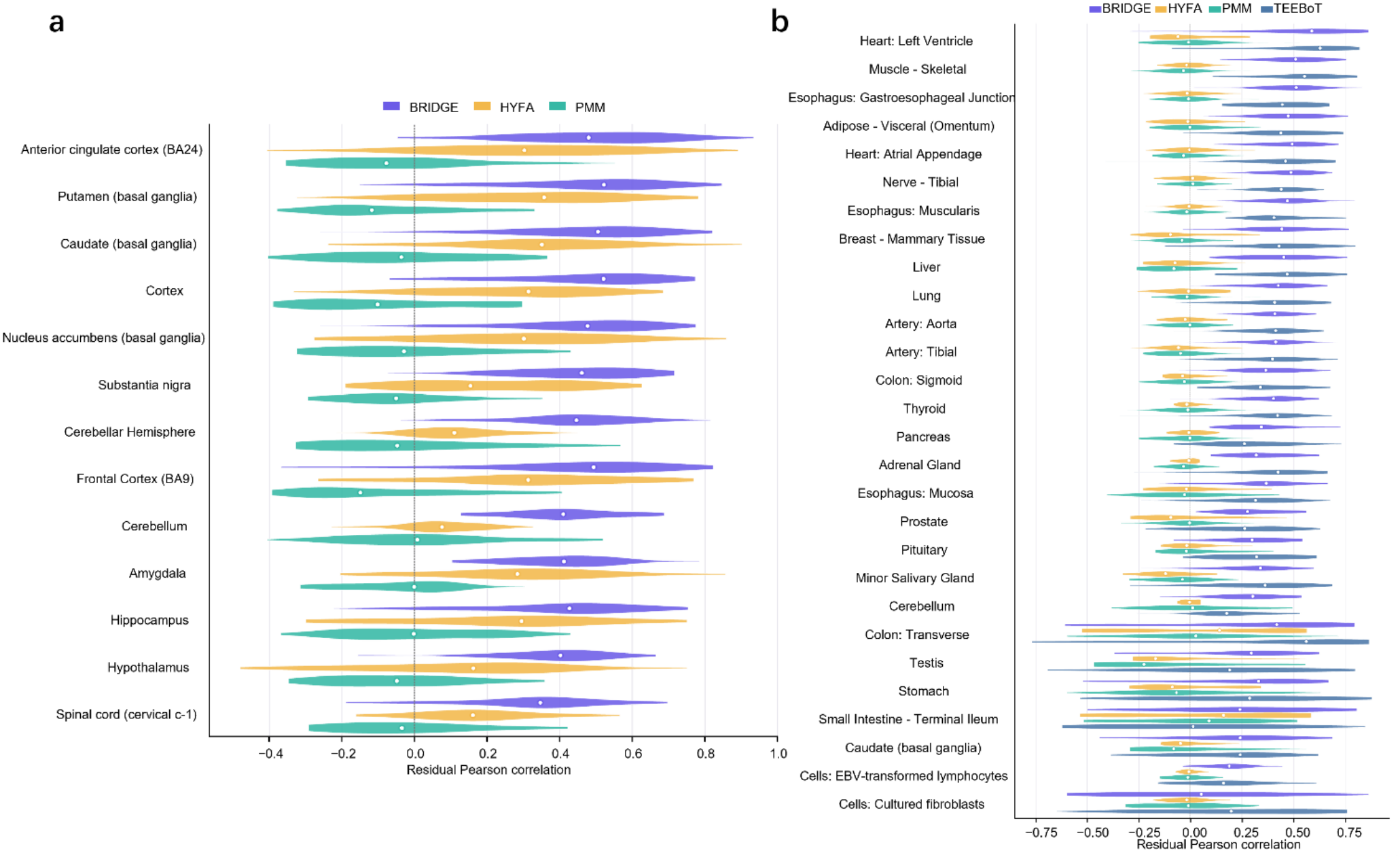
Residual expression recovery in the GTEx hide-one and accessible-tissue evaluations. (a) Distribution of record-level residual Pearson correlations for BRIDGE, HYFA and PMM across brain tissues in the hide-one task. (b) Distribution of residual Pearson correlations for BRIDGE, HYFA, PMM and TEEBoT across target tissues in the accessible-tissue task.

### Component ablation and model behavior

Component ablation showed that the adaptive mixer improved on either residual expert alone. The donor-level average expert alone achieved a mean residual Pearson correlation of 0.361 and an MAE of 0.262, and the sample matching expert alone achieved 0.318 and 0.263. Combining their predictions raised the residual Pearson correlation to 0.434 and lowered the MAE to 0.246 (**Table 1**).

Removing learned target-tissue identity from both the retrieval query and the adaptive mixer reduced the residual Pearson correlation from 0.434 to 0.272 and raised the MAE from 0.246 to 0.282. Also removing the source-tissue identities from the retrieval query gave a residual Pearson correlation of 0.271 and an MAE of 0.282. Age- and sex-based retrieval recovered little donor-specific signal (residual Pearson correlation 0.009, MAE 0.280). Combined with the donor-level average expert, it improved the residual Pearson correlation to 0.367 and lowered the MAE to 0.258, but it remained below the complete BRIDGE model (**Table 1**).

UMAP projections of query embeddings clustered by target-tissue identity and by broader tissue system^37^ (**Figure 3a**). Retrieved records concentrated along the same-tissue diagonal (**Figure 3b**): for 86.2% of target tissues, the most frequently retrieved tissue was the target tissue itself (median self-retrieval fraction, 0.979). Self-retrieval was strongly associated with the number of training donors per tissue (Spearman ρ = 0.82). Among well-represented tissues, retrieval outside the target tissue mainly involved closely related tissues, such as neighboring brain regions (frontal cortex and cortex; cerebellum and cerebellar hemisphere; putamen and caudate) and adjacent parts of the esophagus. By contrast, tissues represented by only five to eight training donors, mostly cell-type-enriched or subregion samples, showed little self-retrieval and retrieved records from unrelated tissues.

**Figure 3.**
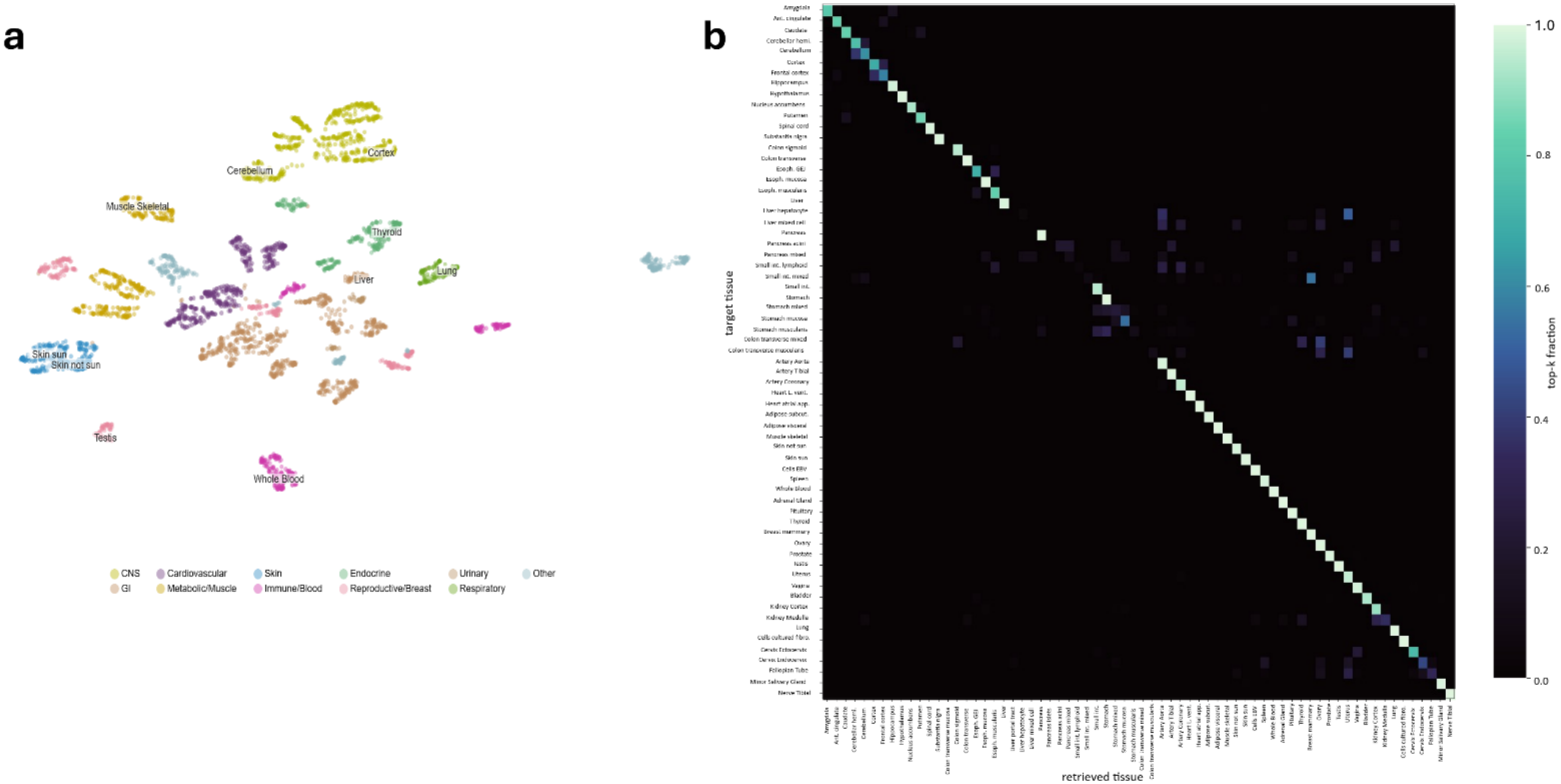
Learned tissue representations and sample matching in BRIDGE. (a) UMAP projection of learned target-query embeddings, colored by broad tissue group. (b) Target-by-retrieved-tissue matrix showing the mean fraction of the top ten memory neighbors drawn from each tissue in the hide-one evaluation.

The adaptive mixer gave the sample matching expert a mean weight of 0.469 across strata defined by the number of observed tissues, with stratum-specific means of 0.456 to 0.484. Weights varied more across target tissues, from 0.291 in kidney cortex to 0.562 in skeletal muscle (**Supplementary Figure 5**).

### BRIDGE improves the recovery of gene–age associations

Across 980 paired evaluations spanning 49 tissues, four missingness levels and 20,000 genes, BRIDGE achieved the highest mean age-β Pearson correlation (0.835), compared with 0.800 for HYFA, 0.744 for target-tissue mean fill, 0.742 for observed-only analysis and 0.370 for PMM. BRIDGE also had a lower age-β MAE than HYFA (0.0231 versus 0.0236) and higher recall of the strongest age-associated genes at the top 100 (0.575 versus 0.505), top 500 (0.612 versus 0.568) and top 1,000 (0.629 versus 0.597) ranks. Sign agreement was similar between the two methods (**Table 2**).

**Table 2.** Recovery of gene-wise age effects after target-tissue masking and reconstruction. BRIDGE was compared with HYFA, PMM, target-tissue mean fill and observed-only analysis across four missingness levels in 49 tissues.

| **Method** | **Age-β Pearson** | **β slope** | **β MAE** | **Sign agreement** | **Top-100 recall** | **Top-500 recall** | **Top-1,000 recall** |
|---|---|---|---|---|---|---|---|
| BRIDGE | 0.835 | 0.649 | 0.0231 | 0.808 | 0.575 | 0.612 | 0.629 |
| HYFA | 0.800 | 0.646 | 0.0236 | 0.810 | 0.505 | 0.568 | 0.597 |
| Mean-fill | 0.744 | 0.488 | 0.0284 | 0.779 | 0.496 | 0.538 | 0.561 |
| Observed-only | 0.742 | 0.945 | 0.0401 | 0.779 | 0.494 | 0.537 | 0.559 |
| PMM | 0.370 | 0.425 | 0.0658 | 0.628 | 0.202 | 0.246 | 0.279 |

Across individual tissues, BRIDGE outperformed HYFA in 794 of 980 paired evaluations (81.0%) and in 43 of 49 tissues after averaging across missingness levels and masks. The highest mean age-β correlations were in transverse colon (0.982), whole blood (0.975), visceral adipose tissue (0.955), esophagus muscularis (0.952) and subcutaneous adipose tissue (0.945). In the six tissues where HYFA performed better, its largest advantage was 0.013 (**Figure 4a**).

**Figure 4.**
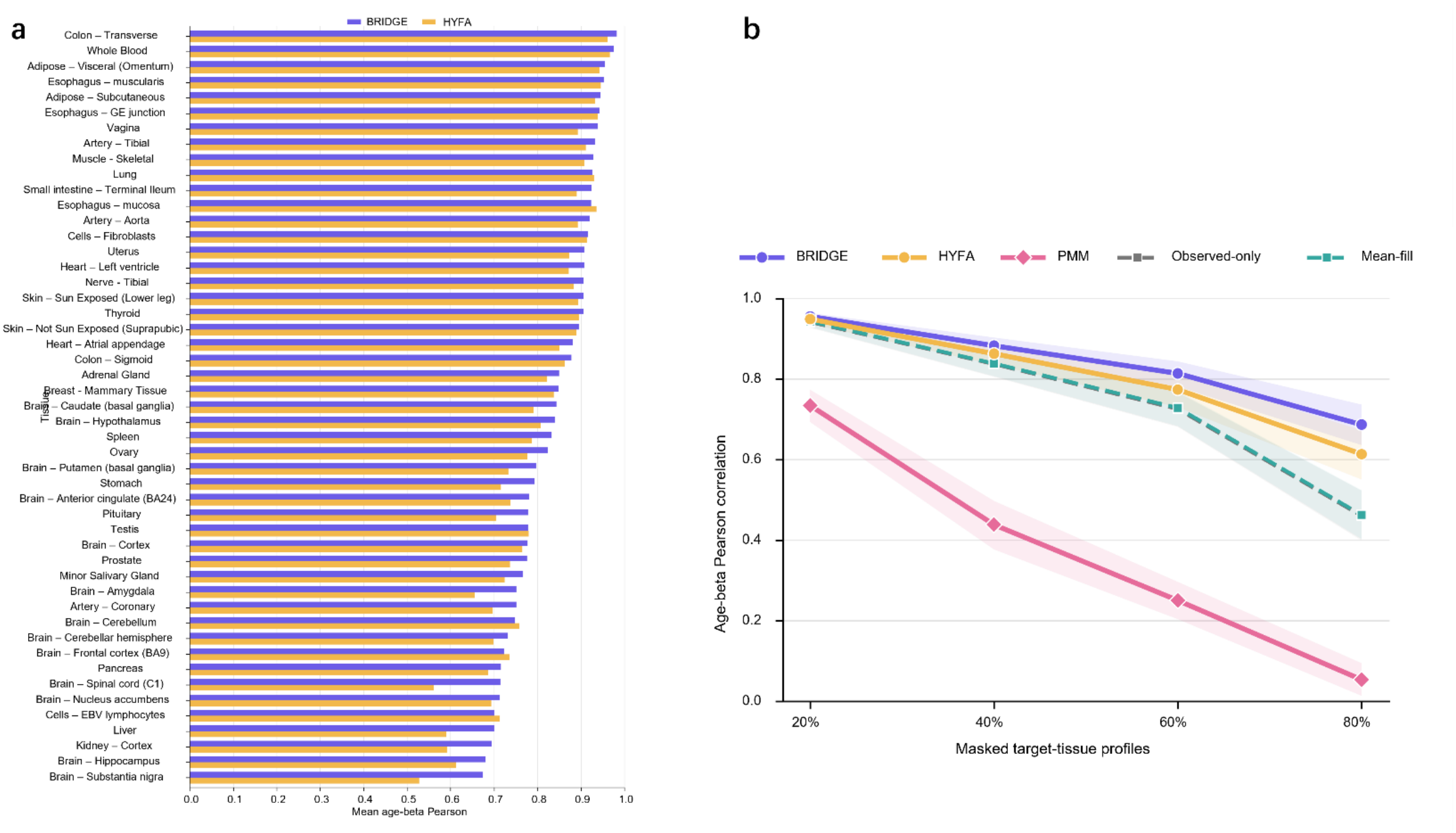
Recovery of gene-wise age effects after target-tissue masking and reconstruction. (a) Mean age-β Pearson correlations for BRIDGE and HYFA across 49 tissues. (b) Mean age-β Pearson correlations across four missingness levels for BRIDGE, HYFA, PMM, observed-only analysis and target-tissue mean fill. Shaded regions indicate 95% bootstrap confidence intervals across tissues.

Age-β recovery declined with increasing missingness for all methods, but BRIDGE had the highest mean correlation at every level: 0.956, 0.883, 0.815 and 0.687 at 20%, 40%, 60% and 80% missingness, respectively, versus 0.949, 0.863, 0.774 and 0.614 for HYFA. The BRIDGE–HYFA advantage therefore grew from 0.007 at 20% missingness to 0.073 at 80% (**Figure 4b**).

### BRIDGE shows Hallmark-specific residual recovery and improvement over HYFA

We next asked whether residual recovery was structured across predefined biological programs. Across 50 MSigDB Hallmark gene sets^34^, BRIDGE achieved a mean residual Pearson correlation of 0.364, with pathway-level values from 0.292 to 0.436 (**Figure 5a, Supplementary Data**). Recovery was highest for TNFα Signaling via NF-κB (0.436), Interferon Alpha Response (0.428) and Interferon Gamma Response (0.425), whereas Pancreas Beta Cells (0.292) and Myogenesis (0.315) were among the least accurately recovered programs (**Figure 5a**). Relative to HYFA, BRIDGE showed significant enrichment of improved genes in 18 of the 50 Hallmark gene sets after Benjamini–Hochberg correction^36^ (FDR < 0.05), with odds ratios from 1.60 to 5.88 (**Figure 5b, Supplementary Data**). The strongest enrichments were for Pancreas Beta Cells (odds ratio = 5.88), Angiogenesis (4.52) and KRAS Signaling Down (4.10). Recovery within selected Hallmark programs also varied across target tissues (**Supplementary Data**).

**Figure 5.**
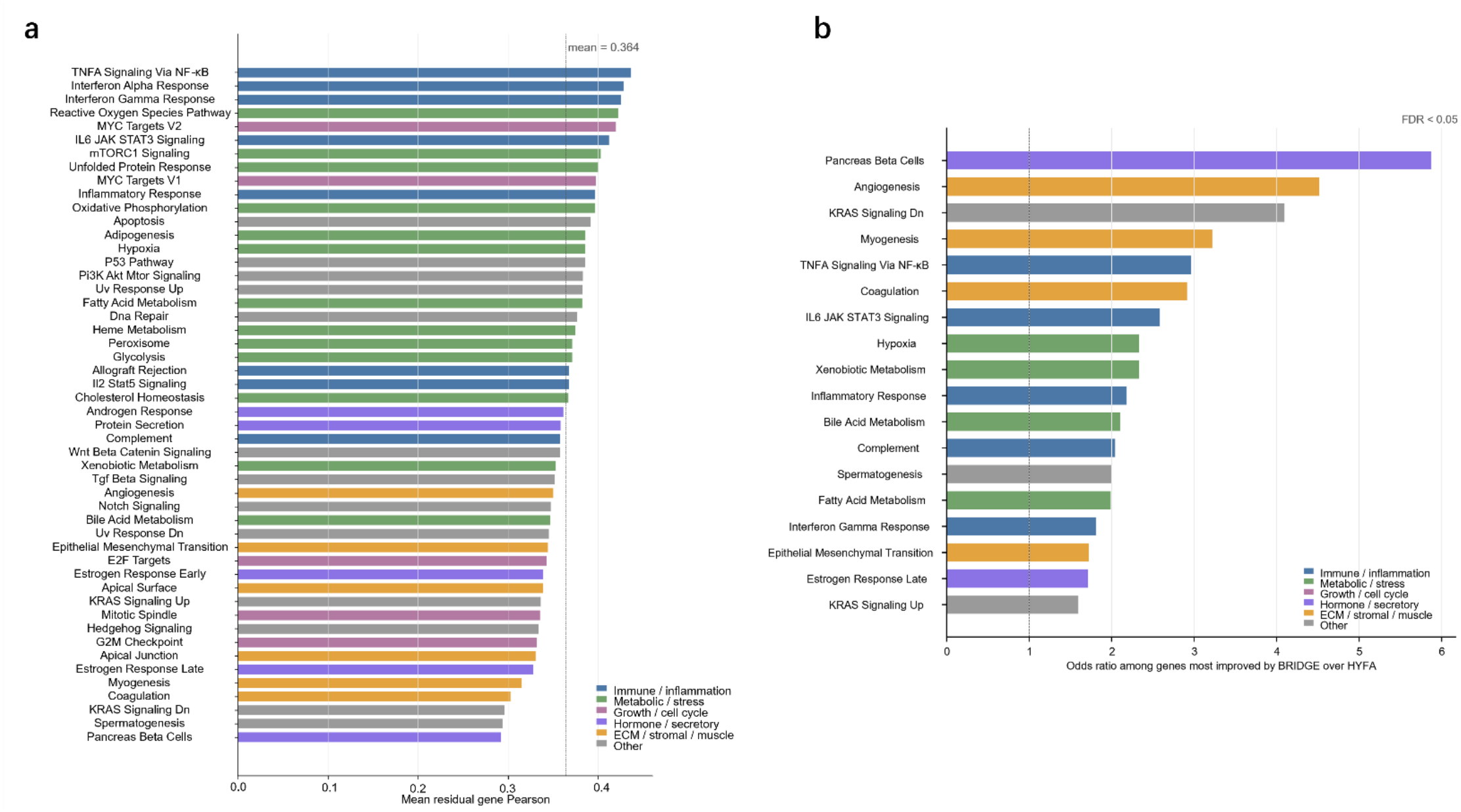
Recovery of Hallmark biological programs by BRIDGE. (a) Mean residual gene Pearson correlation for the 50 MSigDB Hallmark gene sets, ranked by recovery. The dashed line indicates the mean across gene sets. (b) Hallmark gene sets significantly enriched among the top 10% of genes showing the largest MAE improvement with BRIDGE relative to HYFA. The dashed line indicates an odds ratio of 1; only gene sets with FDR < 0.05 are shown. Colors denote biological-program categories.

### BRIDGE converges quickly and benefits from additional training data

Under a shared 200-epoch budget and identical data, BRIDGE reached its best validation MAE (0.2545) at epoch 14, whereas HYFA reached a comparable minimum (0.2554) only at epoch 132. BRIDGE crossed every validation-MAE threshold roughly an order of magnitude earlier; for example, it reached an MAE of 0.260 or lower in 5 epochs versus 72 for HYFA, a 14.4-fold reduction (**Figure 6a,b**).

**Figure 6.**
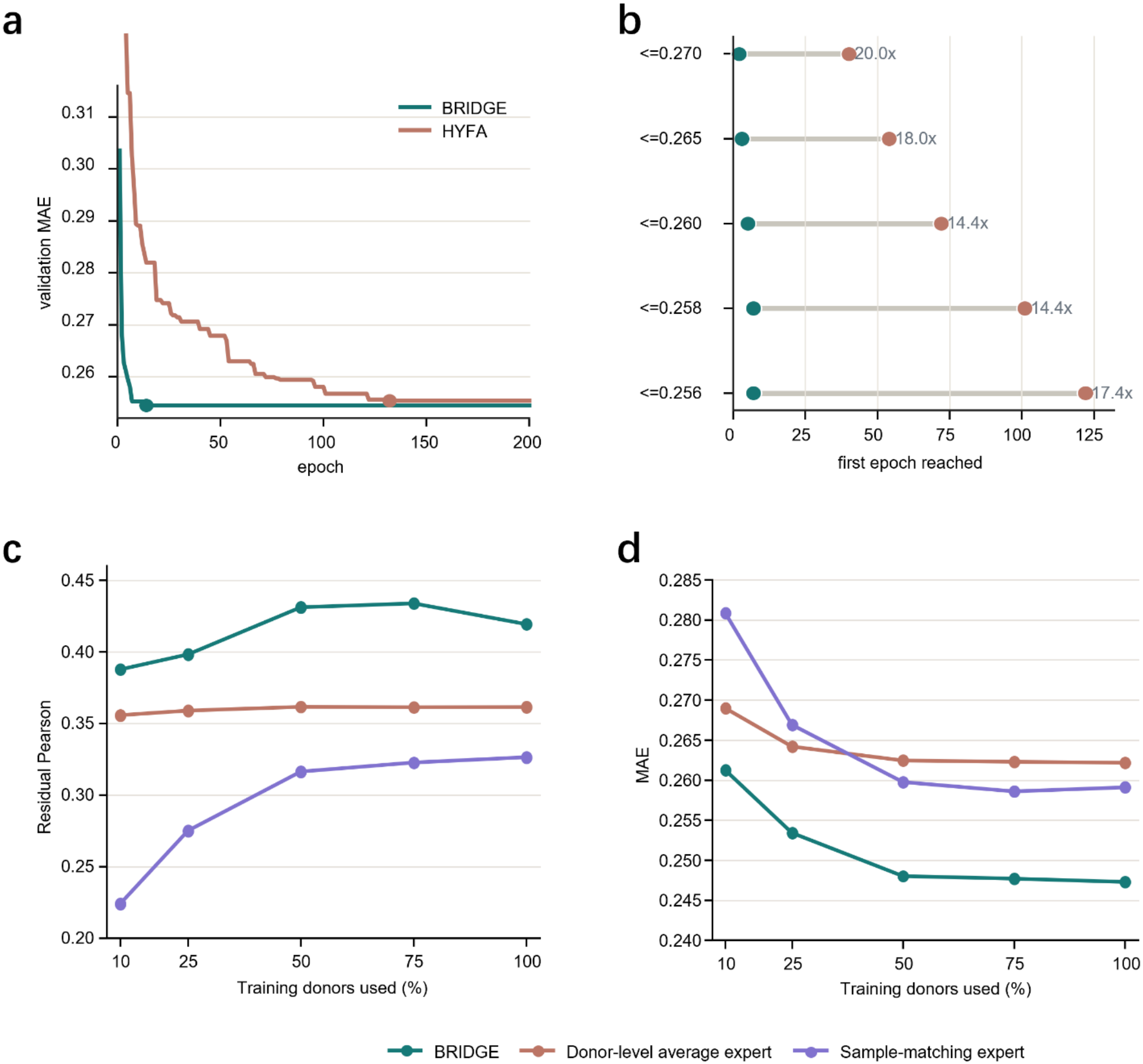
Training efficiency and training-data scaling of BRIDGE. (a) Validation MAE across epochs for BRIDGE and HYFA in the hide-one task. (b) First epoch at which each method reached the indicated validation-MAE threshold. (c) Residual Pearson correlation for BRIDGE and its two component experts using 10%, 25%, 50%, 75% and 100% of the training donors. (d) MAE under the same training-donor fractions.

Retraining BRIDGE on 10–100% of the training donors, with the validation and test sets held fixed, showed that performance improved steadily with more data: residual Pearson correlation increased from 0.388 at 10% to 0.432 at 100%, and MAE decreased from 0.261 to 0.247 (**Figure 6c,d, Supplementary Table 1**). Even with 10% of the training donors, BRIDGE exceeded the residual Pearson correlation of HYFA trained on the full training set (0.237; **Table 1**). The gain was driven by the sample matching expert, whose residual Pearson correlation rose from 0.224 to 0.328 without reaching a plateau, whereas the parameter-free donor-level average expert stayed nearly constant (0.356–0.361). The full model outperformed either expert at every fraction.

### BRIDGE improves recovery on Tabula Sapiens data

To further validate our method, we retrained BRIDGE on the Tabula Sapiens single-cell RNA-seq (scRNA-seq) dataset after pseudobulking the raw counts (**Supplementary Materials**). We found that BRIDGE continued to demonstrate higher residual recovery than HYFA, although this evaluation included only two test donors, due to the smaller sample size of the resource, and should therefore be regarded as preliminary (**Supplementary Table 2**).

### Construction of a BRIDGE-completed GTEx resource

We applied BRIDGE to build a measured-plus-imputed GTEx v11 expression resource spanning 946 donors, 68 tissue labels and 20,000 genes. All 19,788 measured donor–tissue profiles were retained unmodified, and 37,030 additional profiles were imputed across the 60 tissues with sufficient measured support. The remaining eight tissues are kept in the resource metadata but were not imputed. Quality control found no duplicate predictions, no overlap between measured and imputed entries, no query-donor retrieval leakage and no non-finite values. The resulting 10.44-GB chunked HDF5 file contains the measured and imputed expression profiles, the predictions of both residual experts, the mixer weights, gene and tissue identifiers, and a donor-by-tissue mask that distinguishes measured, imputed and non-imputed profiles.

## Discussion

In this study, we introduced BRIDGE, a novel, lightweight, and scalable contrastive learning framework to recover individual-level gene expression from an unobserved target tissue. We compared our method to state-of-the-art techniques such as HYFA and found that BRIDGE was better able to recover complex gene expression profiles in the GTEx dataset, excelling particularly in observed accessible tissues.

Our results have several implications. First, they show that it is insufficient to evaluate imputation methods based on how well they reconstruct the full gene expression profile. The mean profile of the target tissue alone reaches a sample-wise Pearson correlation of 0.969, while carrying no information about the individual donor. The real task is therefore to recover the donor-specific part of expression, meaning how a donor differs from the tissue mean. BRIDGE was designed around this idea. It separates the donor-specific signal into two sources: what the donor’s own observed tissues reveal, and what can be learned from similar donors in a reference population. Each source is modeled by its own expert, and the two are combined according to the information available. With this simple design, and using only tissue identity and gene expression, BRIDGE clearly outperformed HYFA, with the largest gap when only accessible tissues were observed.

Second, our sample matching expert demonstrates the importance of the training objective. A regression model learns to predict the average target given its inputs. When the inputs say little about a donor, that average moves toward the tissue mean and the donor-specific signal is lost. Contrastive learning^23^ asks a different question: which measured target profile belongs to this donor? Because queries and targets are both expressed as differences from the tissue mean, tissue-level expression cannot answer it, so the model must learn features that distinguish donors. The similarity learned this way was far more useful than similarity in age and sex, which recovered almost no donor-specific signal. The retrieval results also show what the model learned and where it falls short. For tissues with enough training donors, retrieval mostly returned records from the same tissue or from closely related tissues, such as neighboring brain regions or different parts of the esophagus. For tissues with only a few donors, retrieval moved to unrelated tissues. Retrieval quality therefore depends on how well each tissue is represented in the reference memory.

The ablation and pathway results show how donor-specific variation behaves. Each expert alone recovered part of the signal, and their combination did better than either, so the two experts capture different information. Removing target-tissue identity caused the largest drop among the ablations, which suggests that a donor’s deviation is not the same in every tissue and must be transferred with the target tissue in mind. The pathway analysis is consistent with this view. Immune, interferon and stress-related programs were recovered best and likely reflect body-wide donor states that appear in many tissues^4^. Programs tied to local tissue structure and function, such as muscle and endocrine programs, were recovered least well. Because GTEx samples are collected after death, some whole-body signals, such as inflammation and hypoxia, may reflect the circumstances of death and tissue collection rather than the donor’s condition during life^38^.

Our age association analysis shows what imputed data can and cannot be used for. BRIDGE’s advantage over HYFA grew as more data were missing, which is the situation most relevant to GTEx. However, both methods shrank age effects relative to those estimated from measured data, by a similar amount, which is a general feature of imputation^32^. Imputed profiles are therefore suitable for finding the direction of effects, ranking genes and selecting candidates, but effect sizes should be corrected before they are interpreted quantitatively, for example with recently developed methods for statistical inference on predicted data^39,40^.

This study has several limitations. Missing tissues were simulated by hiding measured samples, whereas tissues in GTEx are not missing at random^32^. GTEx donors are mostly of European ancestry^2^, and all samples were collected after death, so poorly represented donors may have less reliable neighbors. In addition, most donor-specific variation is still not recovered by BRIDGE, and part of it, such as differences in cell composition between individual samples^42,43^, may not be predictable from other tissues. The next step is to test whether BRIDGE-completed data improve eQTL discovery^5^. This raises its own questions, because the donor-level average expert carries the donor’s own genetic effects, whereas the sample matching expert borrows expression from other donors with different genotypes.

We release a GTEx v11 resource that combines measured and BRIDGE-imputed profiles. The two are kept clearly separate, and each prediction is labeled with the number of observed tissues, quality-control results and expert weights. BRIDGE is also built to improve as GTEx grows. One of its two experts is a memory of measured donor profiles, and the sample matching expert was still improving when trained on the full training set. As new donors are added, the reference becomes larger and more diverse, and because BRIDGE trains quickly, it can be retrained with each new release. More generally, the idea behind BRIDGE is not limited to GTEx: missing measurements for an individual can be predicted by combining that individual’s own data with the experience of similar individuals in a reference population. This approach could be applied to other incomplete multi-tissue cohorts^44^ and, with suitable changes, to other types of incomplete biological data.

## Author contributions

J.Y. conceived the study, collected the data, performed the analysis, developed the model and wrote the first draft of the manuscript. R.Y.L. supervised the work, provided feedback and substantially edited the manuscript. Z.S.Q. supervised the study and provided feedback on the manuscript. All authors reviewed and approved the final manuscript.

## Data availability

GTEx v11 open-access data, including gene expression matrices and sample annotations, are available from the GTEx Portal (https://gtexportal.org). Controlled-access data, including donor genotypes and exact donor ages, are available to approved researchers through dbGaP under accession phs000424.v11.p2, with protected data files hosted on AnVIL. The BRIDGE-completed gene expression tensor, which combines measured and imputed profiles, is available on Zenodo: https://doi.org/10.5281/zenodo.24421141.

## Code availability

The BRIDGE source code is available on GitHub: https://github.com/Jingboyang1105/BRIDGE.

## References

1. Lonsdale, J. et al. The Genotype-Tissue Expression (GTEx) project. Nat. Genet. 45, 580–585 (2013).

2. GTEx Consortium. The GTEx Consortium atlas of genetic regulatory effects across human tissues. Science 369, 1318–1330 (2020).

3. GTEx Consortium. Genetic effects on gene expression across human tissues. Nature 550, 204–213 (2017).

4. Melé, M. et al. The human transcriptome across tissues and individuals. Science 348, 660–665 (2015).

5. Viñas, R. et al. Hypergraph factorization for multi-tissue gene expression imputation. Nat. Mach. Intell. 5, 739–753 (2023).

6. Little, R. J. A. Missing-Data Adjustments in Large Surveys. J. Bus. Econ. Stat. 6, 287–296 (1988).

7. Buuren, S. V. & Groothuis-Oudshoorn, K. mice : Multivariate Imputation by Chained Equations in R. J. Stat. Softw. 45, (2011).

8. Cai, C. et al. Is human blood a good surrogate for brain tissue in transcriptional studies? BMC Genomics 11, 589 (2010).

9. GTEx Consortium et al. Using an atlas of gene regulation across 44 human tissues to inform complex disease- and trait-associated variation. Nat. Genet. 50, 956–967 (2018).

10. Gamazon, E. R. et al. A gene-based association method for mapping traits using reference transcriptome data. Nat. Genet. 47, 1091–1098 (2015).

11. Gusev, A. et al. Integrative approaches for large-scale transcriptome-wide association studies. Nat. Genet. 48, 245–252 (2016).

12. Zhou, D. et al. A unified framework for joint-tissue transcriptome-wide association and Mendelian randomization analysis. Nat. Genet. 52, 1239–1246 (2020).

13. Wang, J. et al. Imputing Gene Expression in Uncollected Tissues Within and Beyond GTEx. Am. J. Hum. Genet. 98, 697–708 (2016).

14. Sul, J. H., Han, B., Ye, C., Choi, T. & Eskin, E. Effectively Identifying eQTLs from Multiple Tissues by Combining Mixed Model and Meta-analytic Approaches. PLoS Genet. 9, e1003491 (2013).

15. Basu, M., Wang, K., Ruppin, E. & Hannenhalli, S. Predicting tissue-specific gene expression from whole blood transcriptome. Sci. Adv. 7, eabd6991 (2021).

16. Xu, W., Liu, X., Leng, F. & Li, W. Blood-based multi-tissue gene expression inference with Bayesian ridge regression. Bioinformatics 36, 3788–3794 (2020).

17. Ghosh, A. J. et al. LungGENIE: the lung gene-expression and network imputation engine. BMC Genomics 26, 227 (2025).

18. Xu, Y. et al. A multi-modal framework improves prediction of tissue-specific gene expression from a surrogate tissue. eBioMedicine 107, (2024).

19. He, G., Chen, M., Bian, Y. & Yang, E. MTM: a multi-task learning framework to predict individualized tissue gene expression profiles. Bioinformatics 39, btad363 (2023).

20. Dong, J., Brown, S. & Truong, K. Estimating genotype-tissue specific gene expression using hybrid deep learning. Commun. Biol. 9, 1019 (2026).

21. Jacobs, R. A., Jordan, M. I., Nowlan, S. J. & Hinton, G. E. Adaptive Mixtures of Local Experts. Neural Comput. 3, 79–87 (1991).

22. Zaheer, M. et al. Deep Sets. Preprint at 10.48550/arXiv.1703.06114 (2018).

23. van den Oord, A., Li, Y. & Vinyals, O. Representation Learning with Contrastive Predictive Coding. ArXiv Prepr. ArXiv180703748 https://doi.org/10.48550/arXiv.1807.03748 (2018) doi:10.48550/arXiv.1807.03748.

24. Radford, A. et al. Learning Transferable Visual Models From Natural Language Supervision. Proc. 38th Int. Conf. Mach. Learn. https://arxiv.org/abs/2103.00020 (2021).

25. Chen, T., Kornblith, S., Norouzi, M. & Hinton, G. A Simple Framework for Contrastive Learning of Visual Representations. Preprint at 10.48550/ARXIV.2002.05709 (2020).

26. Wu, Z., Xiong, Y., Yu, S. & Lin, D. Unsupervised Feature Learning via Non-Parametric Instance-level Discrimination. Preprint at 10.48550/arXiv.1805.01978 (2018).

27. Khandelwal, U., Levy, O., Jurafsky, D., Zettlemoyer, L. & Lewis, M. Generalization through Memorization: Nearest Neighbor Language Models. Preprint at 10.48550/arXiv.1911.00172 (2020).

28. Troyanskaya, O. et al. Missing value estimation methods for DNA microarrays. Bioinformatics 17, 520–525 (2001).

29. Girshick, R. Fast R-CNN. in 2015 IEEE International Conference on Computer Vision (ICCV) 1440–1448 (IEEE, Santiago, Chile, 2015). doi:10.1109/ICCV.2015.169.

30. Jia, K., Cui, C., Gao, Y., Zhou, Y. & Cui, Q. An analysis of aging-related genes derived from the Genotype-Tissue Expression project (GTEx). Cell Death Discov. 4, 91 (2018).

31. Yang, J. et al. Synchronized age-related gene expression changes across multiple tissues in human and the link to complex diseases. Sci. Rep. 5, 15145 (2015).

32. Little, R. & Rubin, D. Statistical Analysis with Missing Data, Third Edition. (Wiley, 2019). doi:10.1002/9781119482260.

33. Subramanian, A. et al. Gene set enrichment analysis: A knowledge-based approach for interpreting genome-wide expression profiles. Proc. Natl. Acad. Sci. 102, 15545–15550 (2005).

34. Liberzon, A. et al. The Molecular Signatures Database Hallmark Gene Set Collection. Cell Syst. 1, 417–425 (2015).

35. Fisher, R. A. On the Interpretation of χ 2 from Contingency Tables, and the Calculation of P. J. R. Stat. Soc. 85, 87 (1922).

36. Benjamini, Y. & Hochberg, Y. Controlling the False Discovery Rate: A Practical and Powerful Approach to Multiple Testing. J. R. Stat. Soc. Ser. B Stat. Methodol. 57, 289–300 (1995).

37. McInnes, L., Healy, J. & Melville, J. UMAP: Uniform Manifold Approximation and Projection for Dimension Reduction. Preprint at 10.48550/arXiv.1802.03426 (2020).

38. Ferreira, P. G. et al. The effects of death and post-mortem cold ischemia on human tissue transcriptomes. Nat. Commun. 9, 490 (2018).

39. Angelopoulos, A. N., Bates, S., Fannjiang, C., Jordan, M. I. & Zrnic, T. Prediction-powered inference. Science 382, 669–674 (2023).

40. Miao, J., Miao, X., Wu, Y., Zhao, J. & Lu, Q. Assumption-lean and data-adaptive post-prediction inference. J. Mach. Learn. Res. 26, 179:9092–179:9122 (2026).

41. Jones, R. C. et al. Tabula Sapiens 2.0: A comprehensive transcriptomic atlas of human cell types. Cell https://doi.org/10.1016/j.cell.2026.08.010 (2026) doi:10.1016/j.cell.2026.08.010.

42. Kim-Hellmuth, S. et al. Cell type–specific genetic regulation of gene expression across human tissues. Science 369, eaaz8528 (2020).

43. Donovan, M. K. R., D’Antonio-Chronowska, A., D’Antonio, M. & Frazer, K. A. Cellular deconvolution of GTEx tissues powers discovery of disease and cell-type associated regulatory variants. Nat. Commun. 11, 955 (2020).

44. Fang, L. et al. The Farm Animal Genotype–Tissue Expression (FarmGTEx) Project. Nat. Genet. 57, 786–796 (2025).

