## Supplementary Materials for "BRIDGE: inter-tissue imputation with contrastive learning improves recovery of donor-specific gene expression"

### GTEx expression processing and donor-level partitioning

GTEx v11 bulk RNA-sequencing expression profiles[1] were analyzed in a fixed 20,000-gene space. Expression values were represented as $\ln(1 + \text{TPM})$. Sample identifiers were used to derive GTEx donor identifiers, and each profile was assigned to its corresponding tissue label. The complete processed dataset contained 19,788 measured profiles from 946 donors across 68 tissue labels. One donor had only a single measured tissue and could not contribute a hide-one record; all remaining donors had at least two measured tissues.

For benchmark experiments, eligible donors were divided into mutually exclusive training, validation and test partitions using seed 42 and proportions of 70%, 10% and 20%, respectively. To preserve the distribution of observed-tissue availability, donors were stratified according to the number of measured tissues using the intervals $[2,4)$, $[4,8)$, $[8,12)$, $[12,20)$ and $[20, \infty)$. This procedure assigned 662 donors to training, 95 to validation and 188 to testing. No expression profile from a donor assigned to one partition was included in another partition.

For each tissue $t$, the reference expression profile $\mu_t$ was calculated using measured profiles from training donors only. If no training profile was available for a tissue label, the global mean across the training profiles was used as a fallback. Tissue means were fixed after partitioning and were used to define residual expression, construct model inputs and calculate residual-level evaluation metrics. Neither validation nor test profiles contributed to the tissue means used in benchmark experiments.

A measured expression profile $\mathrm{x}_{i,t}$ from donor $i$ and tissue $t$ was represented as the sum of the training-derived tissue mean and its residual:

$$x_{i,t} = \mu_t + r_{i,t} \qquad \mathrm{r}_{\mathrm{i,t}} = \mathrm{x}_{i,t} - \mu_t$$

The observed tissue distributions shown in **Supplementary Figure 1** were calculated from the same fixed donor partition. Donor-level coverage was defined as the number of distinct measured tissues available for each donor. Tissue-level coverage was defined as the proportion of donors with a measured profile for each tissue. Age- and sex-distribution summaries were calculated from the public GTEx demographic categories after removal of duplicate donor entries (**Supplementary Figure 2**).

### Sample-matching encoder architecture

The mathematical formulation of BRIDGE is provided in the main **Methods**. Briefly, the sample-matching encoder maps the residual profiles available from a donor and the requested target-tissue identity to a 128-dimensional query representation. The held-out target residual is mapped independently to a 128-dimensional target representation during contrastive training.

The shared expression encoder received a 20,000-dimensional residual-expression vector and comprised a linear transformation from 20,000 to 512 features, layer normalization[2], a Gaussian error linear unit activation (GELU)[3], dropout with probability 0.1, and a final linear transformation to 128 dimensions. The resulting expression representation was normalized to unit $L_2$ length.

Each tissue label was represented by a learned 32-dimensional embedding. For every observed tissue, its normalized residual-expression representation was concatenated with the corresponding tissue embedding. This 160-dimensional vector was passed through a context projection comprising a linear layer from 160 to 128 features, a Gaussian error linear unit activation and a second 128-dimensional linear layer. The resulting evidence vectors were averaged across the observed tissues to obtain a donor state[4].

Records were padded to a maximum of 64 observed tissues. A binary mask identified genuine and padded positions, and only genuine evidence vectors contributed to the masked mean. The donor state was concatenated with the 32-dimensional embedding of the requested target tissue and passed through a query projection with dimensions $160 \rightarrow 128 \rightarrow 128$, with a Gaussian error linear unit activation between the two linear layers. The query representation was then normalized to unit $L_2$ length.

The target residual used during training was encoded by the same expression encoder used for observed residual profiles. Sharing this encoder placed donor queries and measured target residuals in the same representation space. The target residual was used only to construct the positive contrastive target during training and to populate the measured-profile memory bank; it was not available when imputing a missing target tissue.

### Contrastive optimization

The memory encoder was trained with the symmetric InfoNCE objective[5] defined in the main **Methods**, using a temperature of 0.07[6]. Training used the AdamW optimizer[7] with a learning rate of 0.001, weight decay of 0.0001, minibatch size of 256, and a maximum of 200 epochs, with NumPy and PyTorch random-number generators initialized with seed 42. After each epoch, a memory bank was built from the training records, and the validation records were imputed by top-10 retrieval. The encoder checkpoint with the lowest validation sample-matching MAE was retained. Test records were not used for checkpoint selection.

### Memory-bank construction and retrieval

Benchmark memory banks[6] contained training records only, and no test donor contributed any profile to the memory. Retrieval weights were obtained by applying the softmax function to the unscaled cosine similarities of the ten nearest entries; the contrastive temperature was used only during training and was not applied at inference. When retrieval predictions were generated for training records to train the adaptive mixer, the record itself was excluded from its own candidates.

### Adaptive prediction mixer

The adaptive mixer combined the donor-level average and sample-matching predictions. Its inputs were the requested target-tissue identity and the number of observed tissues available for the query donor. Target tissue was represented by a learned eight-dimensional embedding. The observed-tissue count was divided by 64 and concatenated with the tissue embedding. The resulting nine-dimensional feature vector was passed through a $9 \rightarrow 16 \rightarrow 2$ multilayer perceptron with a GELU activation. A softmax over the two outputs produced non-negative expert weights summing to one.

The mixer was trained after the memory encoder had been selected and fixed. Training used Smooth L1 loss[8] between the mixed prediction and measured target expression, AdamW[7] with a learning rate of $1 \times 10^{-3}$, weight decay of $1 \times 10^{-4}$, minibatch size 4,096 and a maximum of 50 epochs. Validation MAE was calculated after every epoch, and the mixer checkpoint with the lowest validation MAE was retained. Neither residual expert was updated during mixer optimization.

### Hide-one record construction

Hide-one records were constructed independently within the training, validation and test donor partitions. For each measured donor–tissue profile, that profile was designated as the target and every other measured profile from the same donor was assigned as observed evidence. The target profile was excluded from its own observed set. Donors with only one measured tissue could not form an eligible record.

Each record stored the donor identifier, target-profile index, encoded target tissue, indices and tissue labels of all observed profiles, and the number of observed tissues. This construction generated 13,879 training records, 1,986 validation records and 3,922 test records. The sum of these records, 19,787, differs from the 19,788 measured profiles because the single-tissue donor could not contribute to a hide-one query.

All benchmark methods were aligned to the same fixed test-record identifiers. Before metric calculation, record order, target-profile identity, target-tissue identity and gene order were checked for exact agreement across the measured reference and method-specific prediction arrays.

### Accessible-tissue record construction

The four widely accessible source tissues[9] were always supplied in a fixed order (Whole Blood; Skin – Sun Exposed (Lower leg); Skin – Not Sun Exposed (Suprapubic); Adipose – Subcutaneous) and were not themselves used as targets. Records were generated separately within each donor partition. After target tissues with fewer than 25 test records were removed[10], the same set of retained target tissues was applied to the training and validation records.

### HYFA comparison

HYFA[10] was implemented using the hypergraph architecture, forward computation, loss function and donor-level source–target sampling procedures supplied in the authors' original source code.

For controlled comparison, HYFA was retrained on the same processed 20,000-gene GTEx matrix and the same seed-42 donor partition used for BRIDGE. The original HYFA study partition was not used. HYFA training was allowed for a maximum of 200 epochs. The formal 20,000-gene run used a minibatch size of 16, and the learning rate used by the matched HYFA configuration. Model parameters were optimized with Adam[11], and the checkpoint with the lowest validation MAE was retained. The reduced minibatch size reflected the memory requirements of the 20,000-gene hypergraph input and was not assumed to be identical to that used in the original HYFA experiments.

The trained HYFA model was evaluated on the fixed hide-one and accessible records rather than on newly sampled evaluation batches. Consequently, BRIDGE and HYFA predicted the same held-out target tissue from the same observed source tissues for every paired comparison. The same record and gene order were enforced before metric calculation.

For the downstream age-association experiment, the primary HYFA prediction set was generated without demographic covariates to prevent age information from entering the imputation model before age-association testing. Demographic-assisted HYFA predictions, where available, were treated only as a sensitivity analysis and were not used for the primary downstream comparison.

### TEEBoT comparison

In the accessible-tissue task, we also evaluated the TEEBoT-style implementation[12] used in the HYFA study. A separate model was fitted for each target tissue using training donors only. For each donor, the expression profiles of the four accessible source tissues were concatenated into a single input vector. The input matrix was centered using the training mean, reduced to 30 components by truncated singular-value decomposition, and used to predict the target-tissue expression profile by multi-output ordinary least-squares regression. In the primary version, age (scaled by 1/100) and sex were added to the input before decomposition; an expression-only version was also evaluated as a sensitivity analysis. Both versions were evaluated on the same 1,270 accessible-tissue test records used for BRIDGE and HYFA.

### Predictive mean matching

Predictive mean matching (PMM)[13] was implemented as a single-imputation comparator in $\ln(1 + \text{TPM})$ expression space using seed 42. PMM used the same donor partition and fixed hide-one or accessible test records used for BRIDGE and HYFA. The reference rows contained measured profiles from the 662 training donors; validation and test donors were excluded from the reference data.

PMM was performed separately by gene. Training reference profiles and the observed profiles defining each test query were combined with a missing cell corresponding to the requested target tissue. Records were assigned to deterministic batches such that a query donor appeared no more than once within a batch. This prevented multiple incompatible missing targets from the same donor from being represented as a single multivariate observation. One imputed value was generated for every requested record–gene combination. The completed formal run covered 20,000 genes, 3,922 hide-one records and 1,270 accessible-tissue records. All returned predictions were required to be finite and to match the requested record, gene, donor and target-tissue keys exactly.

### Component ablations and demographic retrieval

The donor-level average expert and sample-matching expert were evaluated individually by using their predictions before entering the adaptive mixer. The complete BRIDGE prediction used the validation-selected adaptive mixer. Tissue means, donor partitions, test records and evaluation metrics were unchanged across these variants.

The demographic retrieval control used training records with the requested target tissue as candidate profiles. Age was represented as decade ranges instead of exact values, and sex disagreement incurred a fixed distance penalty. For query donor $i$ and candidate donor $j$, the distance was calculated from the absolute scaled age difference plus a sex-mismatch penalty. The formal configuration used an age scale of 10 years, a sex-mismatch penalty of 3, $k = 10$ neighbors, and a retrieval temperature of 1. Candidate distances were converted to softmax weights, and candidate target-tissue residuals were aggregated accordingly. If fewer than ten eligible candidates were available, all available candidates were used. Training-record predictions excluded the identical record.

The demographic retrieval estimate was evaluated by itself and in combination with the Donor-level average expert. The latter combination used a separately trained adaptive mixer with validation-based checkpoint selection. These experiments were controls for whether age and sex matching alone could account for the residual-recovery performance of BRIDGE.

### Analysis of learned model behavior

Three analyses described the learned representation and retrieval behavior of the trained model on the held-out test records. First, the 128-dimensional query representations were projected to two dimensions with UMAP[14] and colored by broad tissue system (**Figure 3a**). Second, for each target tissue, the tissues represented among its top-10 retrieved training records were counted and normalized to retrieval fractions, forming a target-by-retrieved-tissue matrix. The self-retrieval fraction of each target tissue was defined as the fraction of retrieved records from the same tissue, and its relationship with the number of training donors for that tissue was assessed using Spearman correlation. Third, the mixer weight assigned to the sample-matching expert was averaged across records grouped by the number of observed tissues and, separately, by target tissue (**Supplementary Figure 5a-b**). These analyses describe model behavior and were not interpreted as evidence of biological relationships between tissues.

### Evaluation metrics

For record $i$ with target tissue $t$, the observed and predicted residual vectors were

$$ri, t = xi, t - \mu t, \hat{r}i, t = \hat{x}i, t - \mu t$$

The primary record-level metric was Pearson correlation across the 20,000 genes:

$$\mathrm{corr}(\mathbf{r}_{i,t}, \hat{\mathbf{r}}_{i,t}).$$

The reported overall residual Pearson was the arithmetic mean of the finite record-level correlations. Tissue-stratified values were calculated by averaging the record-level correlations within each target tissue. Violin plots displayed the underlying record-level distributions rather than correlations calculated from tissue-averaged profiles.

MAE was calculated over all evaluated record–gene entries in $\ln(1 + \mathrm{TPM})$ expression space:

$$\frac{1}{NG}\sum_{i=1}^{N}\sum_{g=1}^{G}\left|\hat{x},_{i,g} - x_{i,g}\right|,$$

where $N$ is the number of evaluated records and $G = 20{,}000$. Record-level MAE was calculated by averaging absolute error across genes, and tissue-stratified distributions were formed from these record-level values. Because observed and predicted profiles were centered using the same target-tissue mean, subtraction of that mean did not change their absolute difference.

RMSE was calculated over the same record–gene entries as the square root of the mean squared error in ln(1+TPM) space. Sample-wise Pearson correlation was calculated across the 20,000 genes within each record and then averaged across records, whereas gene-wise Pearson correlation was calculated across records within each gene and then averaged across genes. These expression-level metrics were reported for comparison with conventional evaluations[10]; because both are strongly influenced by target-tissue mean expression, they were not treated as primary measures of donor-specific recovery. Residual gene Pearson correlation was calculated across records within each gene using the residual vectors defined above and then averaged across genes.

### Additional analyses and interpretation

#### *Recovery of gene-wise age associations*

The age-association analysis was restricted to hide-one test records with complete public age and sex information. Target tissues were required to contain at least 25 eligible test records. These criteria retained 3,862 records across 49 tissues. Tissue eligibility and the record set were locked before the expanded association analysis.

Public GTEx ages are released as intervals. Each age category was represented by its decade-bin midpoint and then standardized within each tissue using the mean and population standard deviation of the eligible donors. Sex was encoded as a binary covariate according to the GTEx metadata.

For every tissue, missingness levels of 0.2, 0.4, 0.6 and 0.8 were evaluated. Five masks were generated for each tissue and missingness level. The number of masked donors was the rounded product of tissue sample size and the requested missingness proportion. Donors were stratified jointly by encoded age group and sex. Masking quotas were allocated to the strata in proportion to stratum size, with remaining assignments given according to the largest fractional remainders. Donors were sampled without replacement within each stratum. A deterministic seed was derived from seed 42 together with tissue identity, missingness level and mask repeat. The same masks were used for every analysis arm.

The Oracle arm contained all measured target-tissue profiles. The observed-only arm removed masked donors from the association analysis. The mean-fill arm retained all donors but replaced the masked expression profiles with gene-wise means calculated exclusively from the unmasked donors in the same tissue. For BRIDGE, HYFA and PMM, measured profiles were retained for unmasked donors and only the masked profiles were replaced by the corresponding imputed values.

For each gene $g$, an ordinary least squares model was fitted as

$$Y_{i,g} = \alpha_g + \beta_{\text{age},g} A_i + \beta_{\text{sex},g} S_i + \varepsilon_{i,g}$$

where $A_i$ is the standardized age and $S_i$ is encoded sex. The coefficient of standardized age was the effect estimate of interest.

The Oracle model was fitted once per tissue. Every other arm was fitted separately for each tissue, missingness level and mask repeat. The primary recovery endpoint was Pearson correlation across the 20,000 genes between the arm-specific and Oracle age coefficients. The regression slope of reconstructed coefficients on Oracle coefficients was calculated using a first-order least-squares fit. Beta MAE was the mean absolute difference between reconstructed and Oracle age coefficients. Sign agreement was the fraction of genes for which the two coefficients had the same sign.

For top-$k$ recovery, genes were ranked by the absolute magnitude of their age coefficients. The top 100, 500 and 1,000 genes from the Oracle model were compared with the corresponding arm-specific sets. Recall was calculated as the size of the intersection divided by $k$.

### *Hallmark gene-set analysis'*

Human Hallmark gene sets were obtained from MSigDB release 2025.1[15] using gene-symbol annotations. Hallmark symbols were intersected with the symbols represented in the 20,000-gene panel. A gene set required at least five represented genes to be evaluated. All 50 Hallmark sets satisfied the analysis criteria. Genes belonging to multiple sets contributed independently to each applicable set.

For each gene $g$, residual recovery was calculated across hide-one test records:

$$\text{corr}\left(\mathbf{r}_{\cdot,g}, \hat{\mathbf{r}}_{\cdot,g}\right),$$

where the vectors contain the observed and BRIDGE-imputed residual values for that gene across eligible records. Hallmark recovery was the arithmetic mean of the finite gene-level residual correlations among genes represented in the set. This quantity is a within-program summary of gene-wise recovery and was not interpreted as an independent estimate of genome-wide record-level performance.

Gene-level improvement over HYFA was defined as

$$\Delta_g^{\text{MAE}} = \text{MAE}_g^{\text{HYFA}} - \text{MAE}_g^{\text{BRIDGE}}$$

Positive values therefore indicated lower BRIDGE error. The 10% of evaluated genes with the largest $\Delta_g^{\text{MAE}}$ values were designated as the most improved genes. For each Hallmark set, a one-sided Fisher's exact test compared membership in that set with membership in the most-improved group, using the complete mapped 20,000-gene panel as the background universe. $P$ values were adjusted across the 50 Hallmark sets using the Benjamini–Hochberg procedure[16]. Hallmark sets with $q < 0.05$ were included in the enrichment panel.
For the tissue-stratified analysis, gene-level residual correlations were recalculated using only records from each target tissue. Correlations were then averaged within each Hallmark set to form the tissue-by-program matrix. Tissue–gene combinations without sufficient finite variation for Pearson correlation were treated as missing rather than assigned a zero value.

### *Tabula Sapiens pseudobulk validation*

Tabula Sapiens v2[17] single-cell RNA-sequencing data were processed separately by tissue. Only cells generated using the 10x platform were retained. Within each donor–tissue group, raw gene counts were summed across cells. The resulting pseudo-bulk counts were normalized to counts per million,

$$\frac{c_g}{\sum_h c_h} \times 10^6,$$

and transformed as $\ln(1 + \mathrm{CPM})$.

Pseudo-bulk profiles were aligned to the GTEx 20,000-gene order. Genes not measured in Tabula Sapiens were recorded by an observed-gene mask and were excluded from evaluation rather than interpreted as measured zero expression. A total of 19,192 genes were measured in both datasets.

Donors with at least two measured pseudo-bulk tissues were eligible for the hide-one task. The 19 eligible donors were partitioned at the donor level into 16 training, one validation and two test donors, with no donor shared across splits. For each donor, every measured tissue was held out once as the target while its remaining measured tissues served as observed inputs, and a target profile was required to contain at least 100 cells. Target-tissue means were estimated from the training donors only. This produced 324 training, two validation and 15 test hide-one records over the 19,192 shared genes.

BRIDGE and HYFA were retrained entirely within Tabula Sapiens; no GTEx-trained parameters, tissue means or memory entries were reused. BRIDGE followed the architecture and two-stage procedure of the main Methods with dataset-specific tissue embeddings, tissue means, training records and retrieval memory, training the memory encoder for 200 epochs and the adaptive mixer for 50 epochs. HYFA was retrained on the same training donors and 19,192-gene matrix for 200 epochs. Demographic covariates were not used, and checkpoints were selected on the Tabula Sapiens validation donors.

The selected models were evaluated on the same 15 test hide-one records. Performance was summarized across the 19,192 shared genes using residual Pearson correlation, MAE, RMSE, sample-wise Pearson correlation and gene-wise Pearson correlation, following the definitions given above. The Donor-level average expert, sample-matching expert and target-tissue mean baseline were evaluated on the identical records for comparison.

### *Training convergence and data scaling*

BRIDGE and HYFA were each given a maximum training budget of 200 epochs on the same seed-42 GTEx donor partition and 20,000-gene expression matrix. Validation MAE was recorded after every epoch. Each method retained its model-specific optimization and minibatching procedure; the formal convergence runs used minibatch sizes of 256 for BRIDGE and 16 for HYFA.

Convergence was summarized by the epoch with the lowest validation MAE and the first epoch reaching each prespecified MAE threshold. Additional summaries recorded the first epoch falling within 0.001, 0.002 or 0.005 of the method's own minimum validation MAE. These comparisons measure convergence in epochs. Wall-clock duration, GPU-hours and peak memory were not retained for both neural models and were therefore not inferred from file timestamps or from epoch ratios.

Training-size scaling was performed by seeded subsampling of the 662 training donors at fractions of 10%, 25%, 50%, 75% and 100%. Validation and test donors were unchanged. For every fraction, target-tissue means, hide-one training records, model parameters and the retrieval memory were reconstructed using only the selected training donors. Each model was then evaluated using the fixed validation and test partitions. Accuracy was summarized using residual Pearson correlation and MAE.

### *BRIDGE-completed GTEx expression resource*

The BRIDGE-completed GTEx expression tensor was generated only after benchmark development was complete and was treated as a production analysis rather than an additional test of predictive accuracy. BRIDGE was retrained using all 19,788 originally measured profiles from 946 donors and 68 tissue labels. All eligible measured profiles contributed to target-tissue means and the measured-profile memory. The single-tissue donor was retained as measured data but could not provide a hide-one training record, producing 19,787 full-data hide-one records.

The production memory encoder used the standard 20,000-gene architecture, 128-dimensional representations, minibatch size 256, learning rate $1 \times 10^{-3}$ and a fixed 200-epoch duration. Unlike the benchmark model, the full-data production model had no held-out validation partition and no validation-based checkpoint selection. The final 200-epoch encoder was used. The production adaptive mixer was subsequently trained for a fixed 50 epochs on the full-sized hide-one records.

The memory bank contained originally measured profiles only. During retrieval for a missing donor–tissue combination, every memory entry originating from the query donor was excluded before selecting the top ten neighbors. Imputed profiles were never added to the memory bank and were never supplied as observed evidence for later queries.

A preliminary tissue-support audit classified 60 of the 68 tissue labels as eligible for production imputation. The eight lower-support tissues remained available where measured but were not imputed. For every donor and eligible tissue without a measured profile, all originally measured tissues available from that donor were supplied as the query evidence. This procedure produced 37,030 imputed profiles.

This resource contains 19,788 immutable measured profiles and 37,030 imputed profiles across 946 donors and 20,000 genes. A profile manifest records donor identity, target tissue, observed tissue count and mixer weights for every imputed row. The status matrix distinguishes unavailable, measured and imputed donor–tissue combinations.

Production quality control required that: (1) measured and imputed donor–tissue combinations did not overlap; (2) no donor–tissue prediction was duplicated; (3) all expert and final predictions were finite; (4) no retrieved neighbor originated from the query donor; and (5) no imputed profile was used as evidence for another imputation. File hashes were recorded for the measured-expression tensor, encoder checkpoint, mixer checkpoint and released resource.

### Tables and Figures

**Supplementary Data.** This workbook contains three sheets.
Hide-one. Per-tissue imputation performance on the hide-one task (65 tissues; BRIDGE, HYFA and PMM). Mean residual Pearson correlation and MAE for each method in each target tissue, together with the number of training donors and test records per tissue.
Accessible_tissues. Per-tissue imputation performance on the hide-one task (28 tissues; BRIDGE, TEEBoT, HYFA, and PMM). Mean residual Pearson correlation and MAE for each method in each target tissue, together with the number of training donors and test records per tissue.
Hallmark_recovery. Hallmark-level residual recovery and enrichment of BRIDGE improvements. All 50 MSigDB Hallmark gene sets, ranked by BRIDGE's mean residual gene-wise Pearson correlation, together with the represented genes, accuracy metrics and the mean reduction in MAE relative to HYFA.

**Supplementary Table 1.** Effect of training-set size on BRIDGE and its two experts. BRIDGE was retrained using 10–100% of the 662 GTEx training donors, with the validation and test partitions held fixed. All models were evaluated on the same 3,922 hide-one test records across 20,000 genes. The full model was retrained independently, so its values differ slightly from Table 1.

| Training fraction | Donor-level average expert MAE | Donor-level average expert Residual Pearson | Sample-matching expert MAE | Sample-matching expert Residual Pearson | BRIDGE MAE | BRIDGE Residual Pearson |
|---|---|---|---|---|---|---|
| 10% | 0.2690 | 0.3557 | 0.2808 | 0.2239 | 0.2612 | 0.3877 |
| 25% | 0.2642 | 0.3589 | 0.2682 | 0.2774 | 0.2527 | 0.4072 |
| 50% | 0.2625 | 0.3616 | 0.2607 | 0.3070 | 0.2482 | 0.4241 |
| 75% | 0.2623 | 0.3613 | 0.2620 | 0.3190 | 0.2480 | 0.4262 |
| 100% | 0.2622 | 0.3614 | 0.2598 | 0.3281 | 0.2472 | 0.4321 |

**Supplementary Table 2.** Imputation performance on Tabula Sapiens after within-dataset retraining. BRIDGE and HYFA were retrained on Tabula Sapiens pseudo-bulk profiles (16 training, 1 validation and 2 test donors) and evaluated on the same 15 hide-one test records across 19,192 genes shared with the GTEx panel. MAE and RMSE were calculated in $\ln(1 + \mathrm{CPM})$ space. NA, not applicable.

| Method | Residual Pearson | MAE | RMSE | Sample Pearson | Gene Pearson |
|---|---|---|---|---|---|
| BRIDGE | 0.178 | 0.377 | 0.639 | 0.934 | 0.391 |
| Donor-level average expert | 0.165 | 0.388 | 0.630 | 0.932 | 0.421 |
| Sample-matching expert | 0.122 | 0.410 | 0.696 | 0.924 | 0.351 |
| HYFA | 0.101 | 0.374 | 0.639 | 0.933 | 0.489 |
| Target-tissue mean | NA | 0.350 | 0.605 | 0.938 | 0.521 |

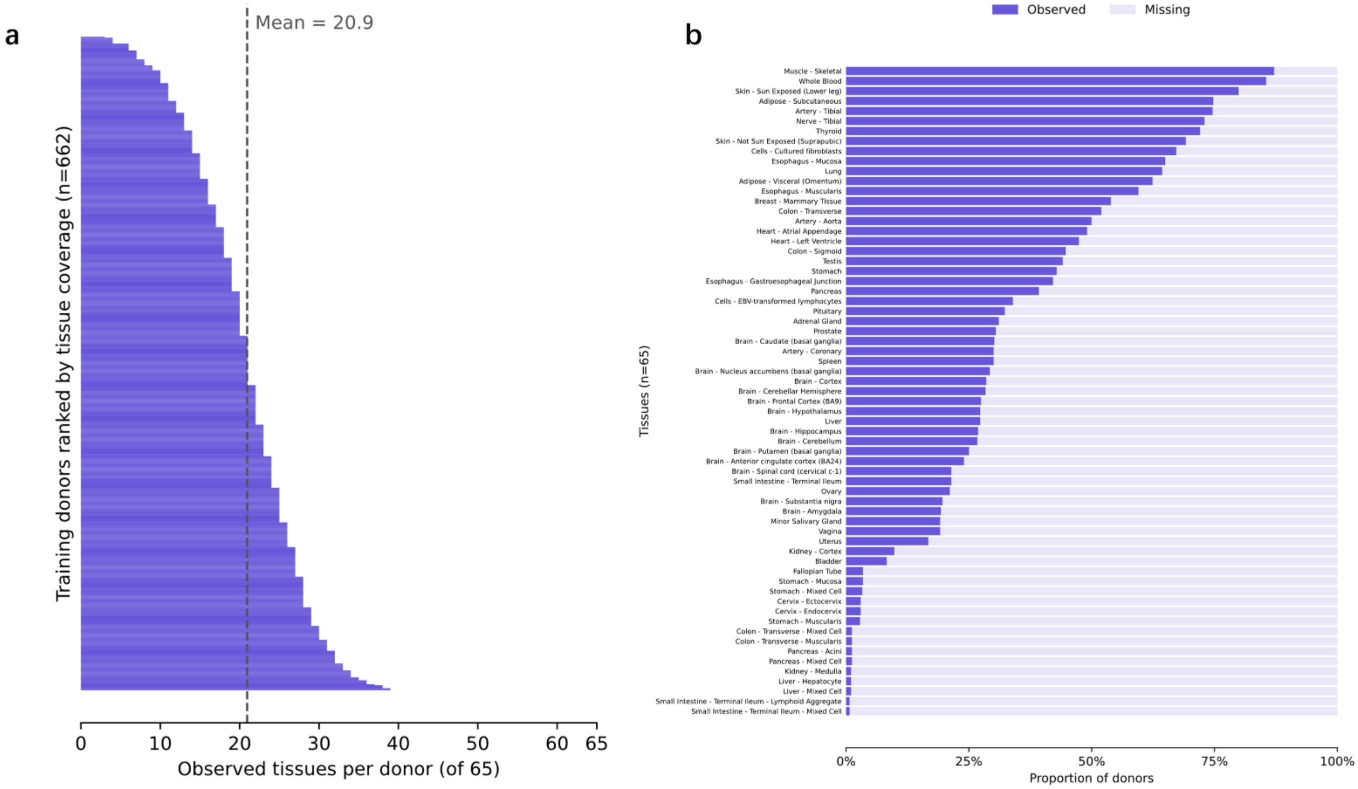


**Supplementary Figure 1.** Tissue availability among GTEx v11 training donors. (a) Number of observed tissues per training donor (n = 662), out of 65 tissues. Donors are ranked by tissue coverage; the dashed line marks the mean (20.9 tissues). (b) Proportion of training donors with observed or missing expression for each of the 65 tissues, ordered by decreasing availability.

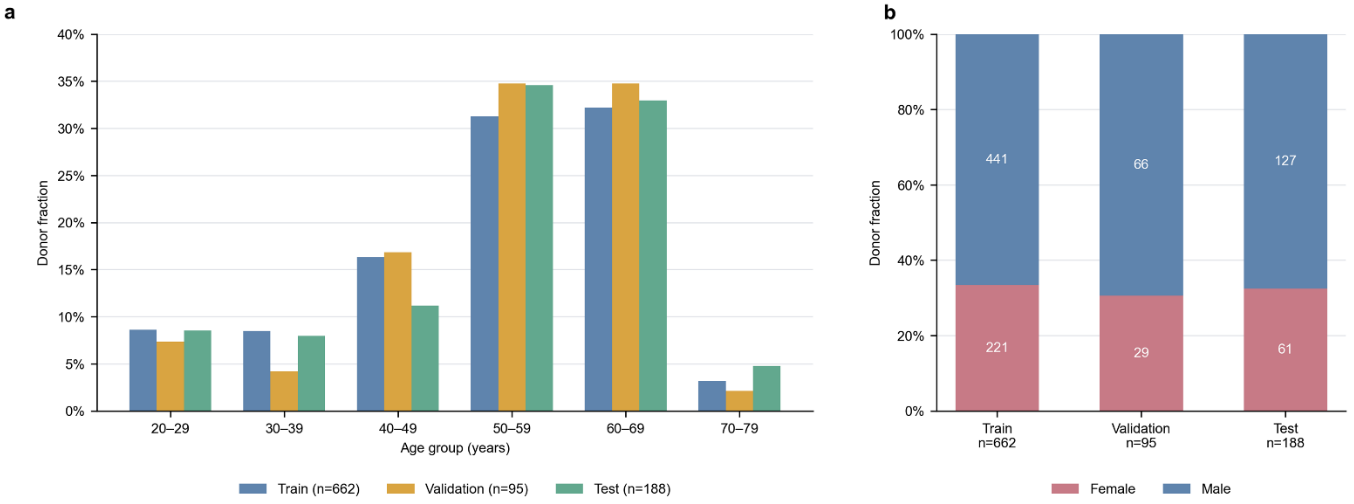


**Supplementary Figure 2.** Age and sex composition of the donor partitions. (a) Distribution of donors across age groups in the training (n = 662), validation (n = 95) and test (n = 188) partitions. (b) Sex composition of each partition, with donor numbers shown within each bar.

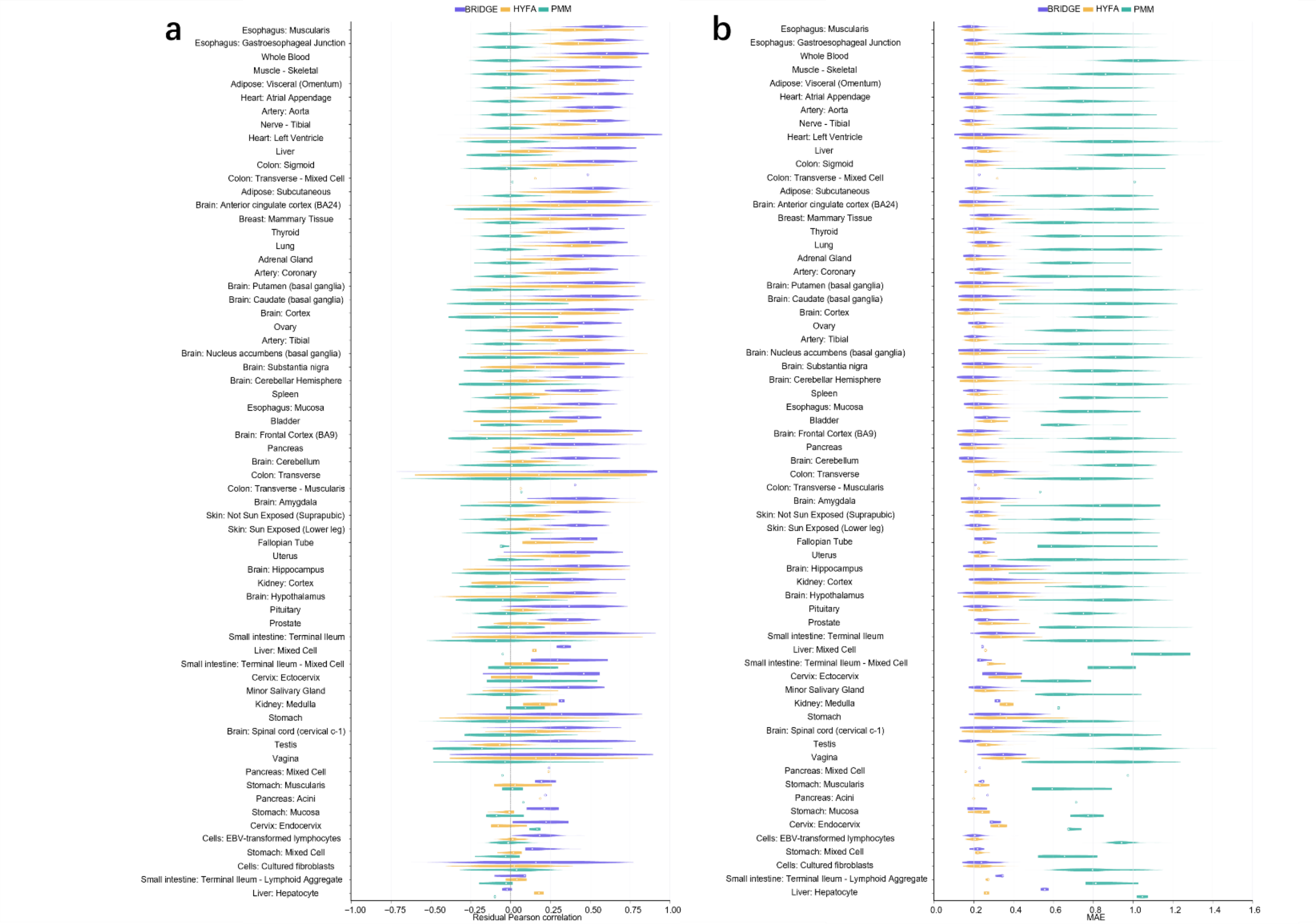


**Supplementary Figure 3.** Hide-one imputation performance across all 65 GTEx tissues. (a) Record-level residual Pearson correlations for BRIDGE, HYFA and PMM in each target tissue. (b) Record-level MAE for the same tissues and records. Tissues are ordered by decreasing mean residual Pearson correlation of BRIDGE, and the same order is used in both panels. Violins show distributions across the 3,922 hide-one test records, and white points indicate medians.

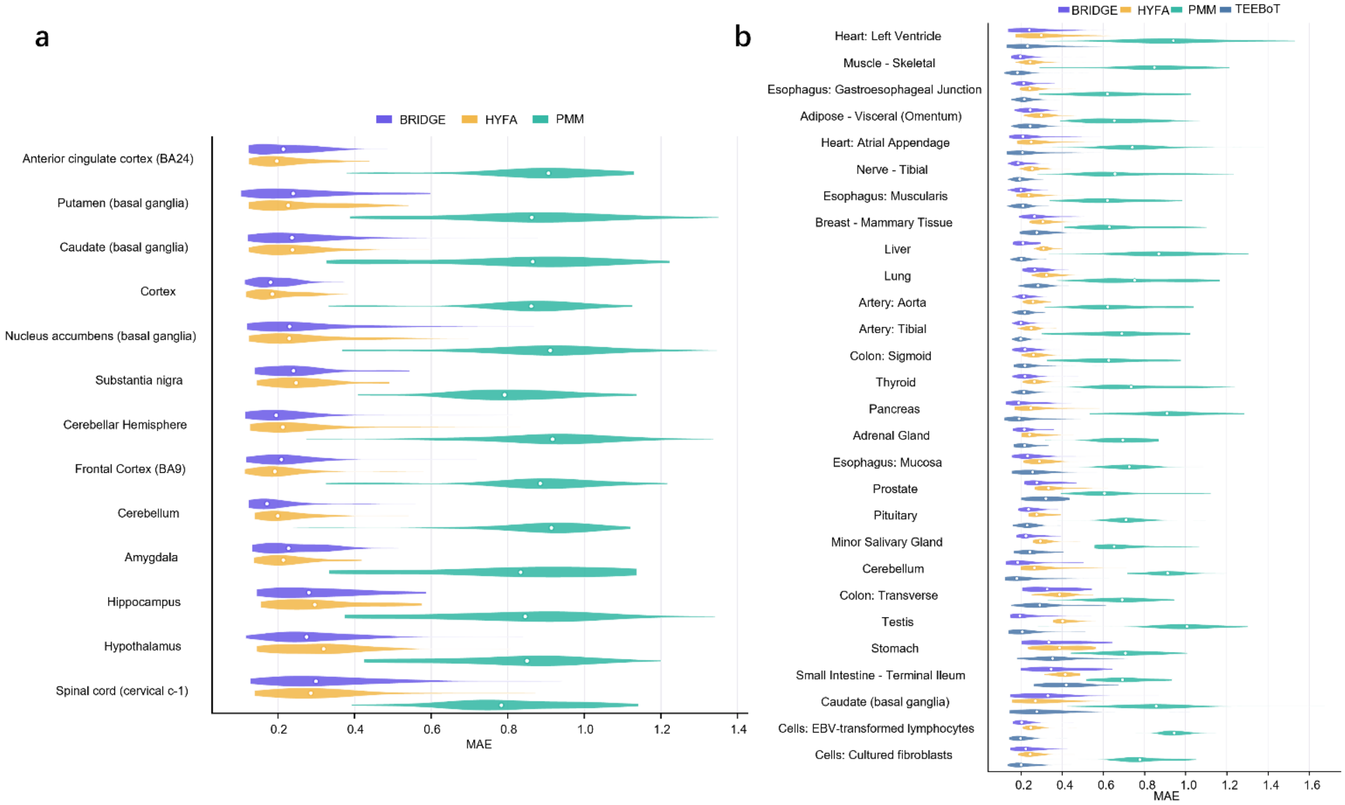


**Supplementary Figure 4.** Mean absolute error of all methods in brain tissues and in the accessible-tissue task. (a) Record-level MAE for BRIDGE, HYFA and PMM across the 13 brain tissues in the hide-one task, complementing the residual Pearson correlations shown in Fig. 2a. (b) Record-level MAE for BRIDGE, HYFA, PMM and TEEBoT across the 28 target tissues in the accessible-tissue task. White points indicate medians.

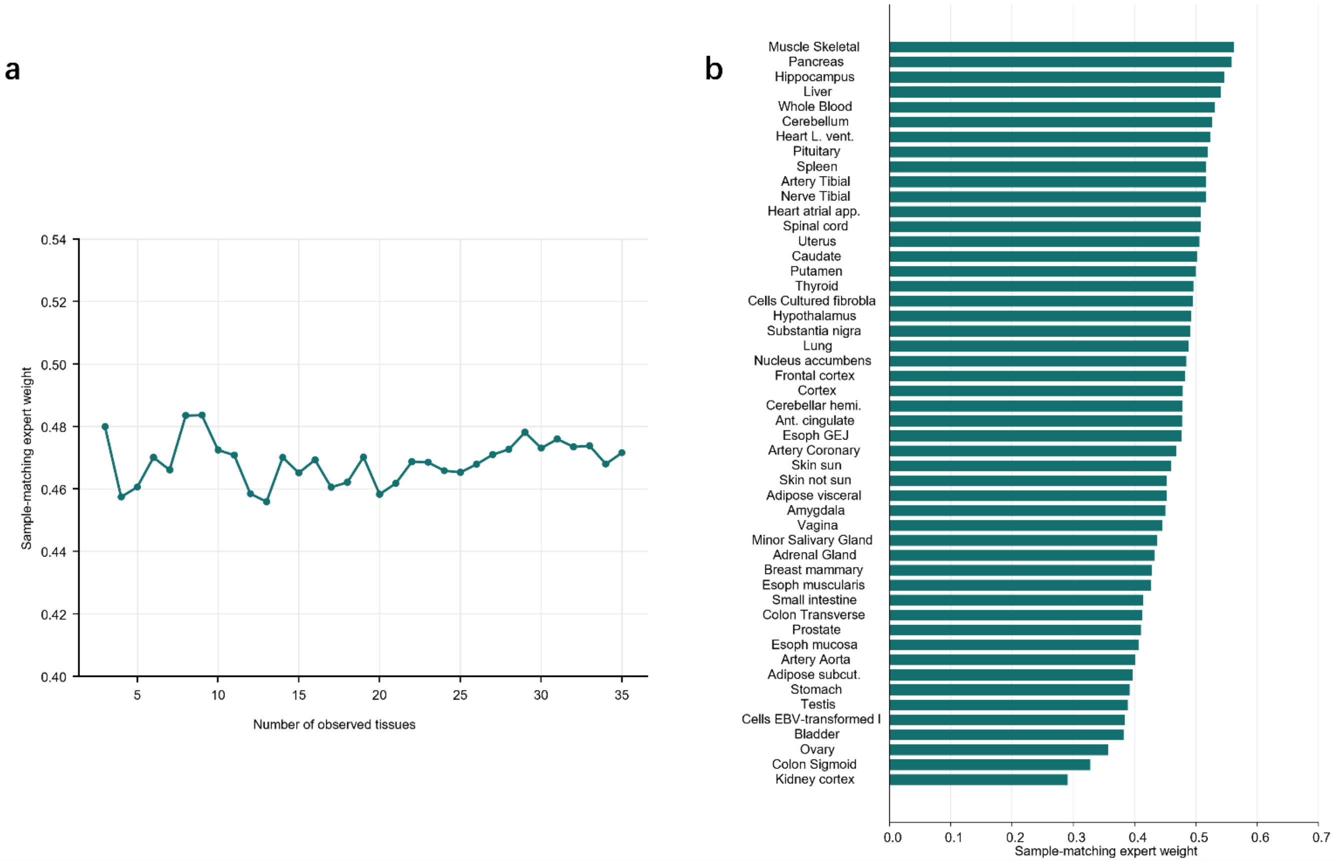


**Supplementary Figure 5.** Weights assigned by the adaptive mixer to the sample-matching expert. (a) Mean sample-matching expert weight as a function of the number of observed tissues per donor. Each point shows the mean across hide-one test records with that number of observed tissues. (b) Mean sample-matching expert weight for each of the 50 target tissues with at least 10 test records, ranked from highest to lowest. The donor-level average expert receives the remaining weight (1 minus the value shown). These panels describe model behavior and are not intended to indicate biological relationships between tissues.

### References


1. Lonsdale, J. *et al.* The Genotype-Tissue Expression (GTEx) project. *Nat. Genet.* **45**, 580–585 (2013).
2. Ba, J. L., Kiros, J. R. & Hinton, G. E. Layer Normalization. Preprint at https://doi.org/10.48550/ARXIV.1607.06450 (2016).
3. Hendrycks, D. & Gimpel, K. Gaussian Error Linear Units (GELUs). Preprint at https://doi.org/10.48550/ARXIV.1606.08415 (2016).
4. Zaheer, M. *et al.* Deep Sets. Preprint at https://doi.org/10.48550/arXiv.1703.06114 (2018).
5. van den Oord, A., Li, Y. & Vinyals, O. Representation Learning with Contrastive Predictive Coding. *ArXiv Prepr. ArXiv180703748* https://doi.org/10.48550/arXiv.1807.03748 (2018) doi:10.48550/arXiv.1807.03748.
6. Wu, Z., Xiong, Y., Yu, S. & Lin, D. Unsupervised Feature Learning via Non-Parametric Instance-level Discrimination. Preprint at https://doi.org/10.48550/arXiv.1805.01978 (2018).
7. Loshchilov, I. & Hutter, F. Decoupled Weight Decay Regularization. Preprint at https://doi.org/10.48550/ARXIV.1711.05101 (2017).
8. Jacobs, R. A., Jordan, M. I., Nowlan, S. J. & Hinton, G. E. Adaptive Mixtures of Local Experts. *Neural Comput.* **3**, 79–87 (1991).
9. Viñas, R. *et al.* Hypergraph factorization for multi-tissue gene expression imputation. *Nat. Mach. Intell.* **5**, 739–753 (2023).
10. Viñas, R. *et al.* Hypergraph factorisation for multi-tissue gene expression imputation. 2022.07.31.502211 Preprint at https://doi.org/10.1101/2022.07.31.502211 (2022).
11. Kingma, D. P. & Ba, J. Adam: A Method for Stochastic Optimization. Preprint at https://doi.org/10.48550/arXiv.1412.6980 (2017).
12. Basu, M., Wang, K., Ruppin, E. & Hannenhalli, S. Predicting tissue-specific gene expression from whole blood transcriptome. *Sci. Adv.* **7**, eabd6991 (2021).
13. Little, R. J. A. Missing-Data Adjustments in Large Surveys. *J. Bus. Econ. Stat.* **6**, 287–296 (1988).
14. McInnes, L., Healy, J. & Melville, J. UMAP: Uniform Manifold Approximation and Projection for Dimension Reduction. Preprint at https://doi.org/10.48550/arXiv.1802.03426 (2020).
15. Melé, M. *et al.* The human transcriptome across tissues and individuals. *Science* **348**, 660–665 (2015).
16. Benjamini, Y. & Hochberg, Y. Controlling the False Discovery Rate: A Practical and Powerful Approach to Multiple Testing. *J. R. Stat. Soc. Ser. B Stat. Methodol.* **57**, 289–300 (1995).
17. Jones, R. C. *et al.* Tabula Sapiens 2.0: A comprehensive transcriptomic atlas of human cell types. *Cell* https://doi.org/10.1016/j.cell.2026.08.010 (2026) doi:10.1016/j.cell.2026.08.010.
